# Dynamics of White Matter Architecture in Lexical Production among Middle-Aged Adults

**DOI:** 10.1101/2024.02.09.579645

**Authors:** Clément Guichet, Élise Roger, Arnaud Attyé, Sophie Achard, Martial Mermillod, Monica Baciu

## Abstract

This study aimed to elucidate the white matter changes associated with lexical production (LP) difficulties that typically emerge in middle age, resulting in increased naming latencies. To delay the onset of LP decline, middle-aged adults may rely on domain-general (DG) and language-specific (LS) compensatory mechanisms as proposed by the LARA model (Lexical Access and Retrieval in Aging). However, our knowledge of the white matter changes supporting these mechanisms remains incomplete. Based on a sample of 155 middle-aged adults from the CAMCAN cohort, we combined dimensionality reduction techniques with multivariate statistical methods to jointly examine the relationships between diffusion-weighted imaging and LP-related neuropsychological data. Our findings (i) show that midlife constitutes a pivotal period marked by a discontinuity in brain structure within distributed networks within dorsal, ventral, and anterior cortico-subcortical pathways, and (ii) reveal that this discontinuity signals a neurocognitive transition around age 53-54, marking the onset of LP decline. Indeed, our results propose that middle-aged adults may initially adopt a “semantic strategy” to compensate for initial LP challenges. Still, this strategy may be compromised when late middle-aged adults (age 55-60) lose the ability to exert cognitive control over semantic representations (i.e., reduced semantic control).

In summary, our study advances our comprehension of brain structure changes that underpin the neurocognitive profile of LP in middle age. Specifically, we underscore the importance of considering the interplay between DG and LS processes when studying the trajectory of LP performance in healthy aging. Furthermore, these findings offer valuable insights into identifying predictive biomarkers related to the compensatory dynamics observed in midlife, which can help understand language-related neurodegenerative pathologies.

**Highlights:**

- Midlife constitutes a pivotal period characterized by a discontinuity in brain structure.
- Early middle-aged adults (age 45-55) adopt a “semantic strategy” to facilitate semantic access and sustain lexical production (LP) performances.
- Late middle-aged adults (age 55-60) gradually lose the ability to exert cognitive control over semantic representations, marking the onset of LP decline.

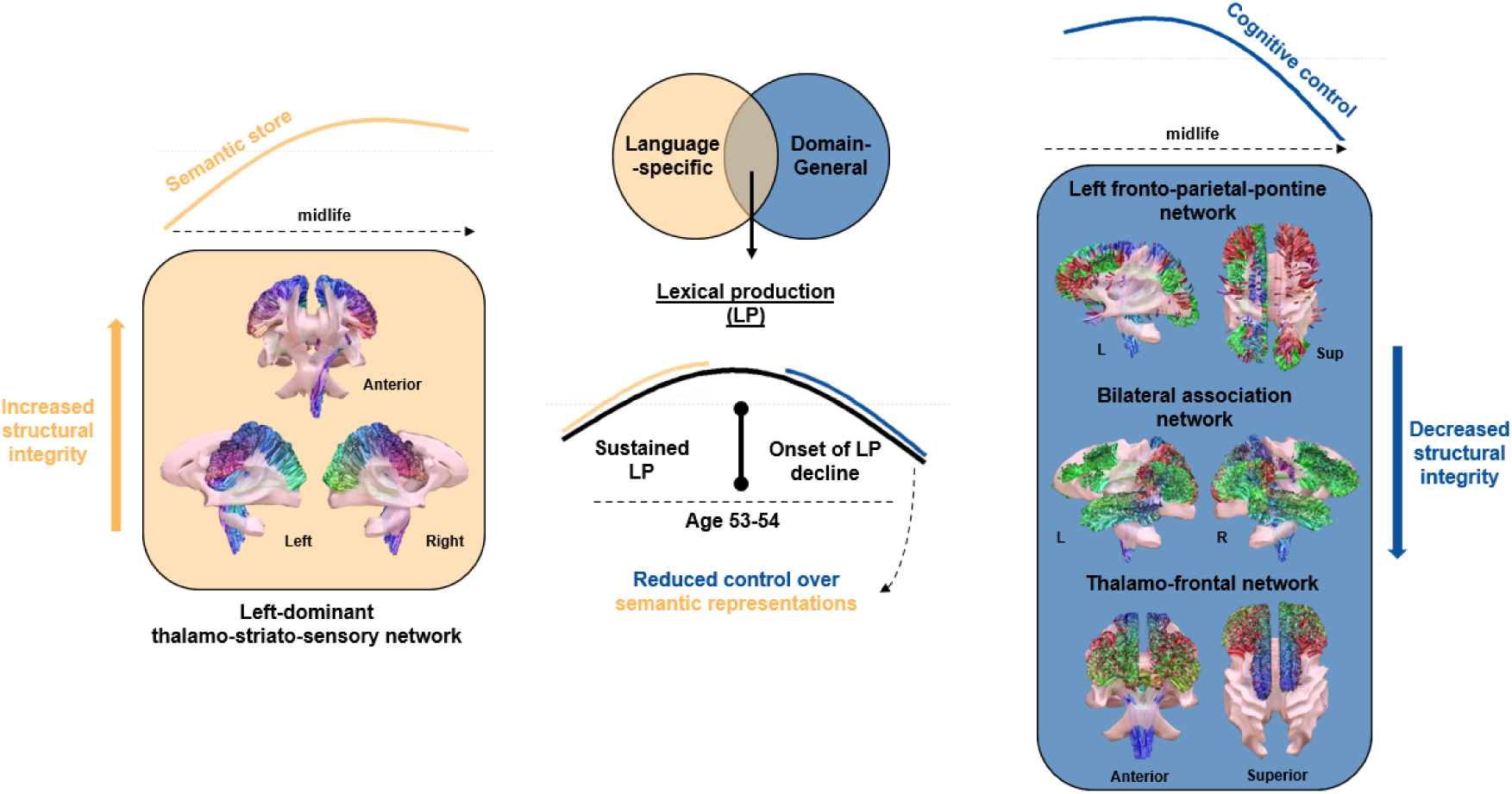

## 1. Introduction

Population aging has led to an increase in average life expectancy. However, this demographic transition is accompanied by a higher prevalence of neurodegenerative diseases and cognitive decline, leading to an increase in healthcare demand (United Nations, 2023). Language function, crucial for human communication and cognition (Hagoort, 2019, 2023), undergoes significant and differentiated changes related to individuals age (De Zubicaray & Schiller, 2018). Indeed, while some aspects of language processes, such as language comprehension (LC), remain relatively stable compared to younger adults, others, such as language production (LP), decline relatively early with age, starting in middle age (Oosterhuis et al., 2023; Verhaegen & Poncelet, 2013). The disparity between LP and LC can be attributed to the benefits from rich semantic knowledge accumulated over the lifespan (Gollan & Goldrick, 2019; Salthouse, 2019). Indeed, this semantic expertise (Spreng & Turner, 2019) contributes to preserving language comprehension and implementing compensatory mechanisms to mitigate LP decline in older adults (Baciu et al., 2021). At the behavioral level, while advanced age is associated with the most significant LP decline (e.g., lower naming accuracy) (Shafto & Tyler, 2014; Verhaegen & Poncelet, 2013), increased naming latencies in low-frequency words with poorer semantic connections may onset in middle age (Benítez-Burraco & Ivanova, 2023). Therefore, a comprehensive investigation of midlife is crucial to understanding the aging mechanisms at play and predicting the neurocognitive trajectory in older late adulthood (Lachman, 2015; Park & Festini, 2016).

Previous studies have reported a gradual LP decline resulting from the interplay between domain-general and language-specific mechanisms (Gertel et al., 2020; Shafto & Tyler, 2014; Steinberg Lowe & Buchwald, 2023). Typically, there is reduced access to and/or retrieval of lexico-semantic representations (Baciu et al., 2016; Boudiaf et al., 2018) in association with a decline in executive functioning with age (Higby et al., 2019), reduced working memory (Burke & Shafto, 2004) and inhibitory processes (Gordon & Kurczek, 2014) explaining the slowdown of information processing and reduced verbal fluency (Gordon et al., 2018; Whiteside et al., 2016) within the LP decline context.

The decline in LP varies among older individuals, exhibiting significant variability (Gordon et al., 2018; Wen & Dong, 2023) linked to cognitive reserve differences (Oosterhuis et al., 2023; Perneczky, 2019). As highlighted by previous studies (Baciu et al., 2021; Wulff et al., 2022) this variability depends on various cognitive reserve factors such as education, social, intellectual, physical, and leisure activities (Brosnan et al., 2023; Perneczky, 2019). As previously reported by the STAC (Scaffolding Theory of Aging and Cognition) model (Park & Reuter-Lorenz, 2009), scaffolding is a natural process throughout life that entails utilizing and enhancing alternative neural pathways to accomplish specific cognitive tasks. Its effectiveness is enhanced by cognitive engagement, physical exercise, and reduced default network activity. Subsequently, a revised version of this model entitled STAC-r (Reuter-Lorenz & Park, 2014, 2023) has been described as incorporating life-course factors that contribute to either bolstering or depleting neural resources, thereby influencing the developmental trajectory of brain structure, function, and cognitive abilities. The life-course factors also impact individuals’ compensatory mechanisms to address cognitive challenges and mitigate the adverse effects of structural and functional decline. Specifically, studies point to midlife lifestyle activities as a key contributor to this idiosyncratic nature of cognitive aging (Chan et al., 2018; Xu et al., 2019). The intricate landscape of language aging underscores the need to comprehend behavioral and cognitive changes and the functional and structural network reorganization associated with compensatory mechanisms. This understanding aims to delay, as much as possible, a decline in LP performance and maintain a relatively normal level.

At a functional level, data from resting-state and task-activation fMRI suggest that aging influences activation dynamics within large brain networks, reorganizing them to maintain cognitive performance (Deery et al., 2023; Sun et al., 2023). With aging, cerebral networks exhibit reduced functional specialization, as seen by decreased within-network connectivity (Chan et al., 2014; Edde et al., 2021; Jockwitz et al., 2017). Simultaneously, increased between-network connectivity may counteract cognitive decline (Bertolero et al., 2015; Cabeza et al., 2018; Meunier et al., 2009). Notably, reduced activity within the Default Mode Network (DMN), primarily involved in semantic processes (Alves et al., 2019; Menon, 2023; Raichle et al., 2001), coupled with enhanced functional integration in control-executive regions play a crucial compensatory role in tasks with significant attentional demands (see the DECHA model; Spreng & Turner, 2019) such as LP (Roger, Banjac, et al., 2022). This aligns with studies showing that semantic cognition becomes increasingly reliant on domain-general processes with age to ensure efficient and context-relevant use of semantic representations (Hoffman & Morcom, 2018; Martin, Saur, et al., 2022; Martin, Williams, et al., 2022). For example, Hoyau et al. (2018) found that, compared to younger adults, older adults modulate the top-down connectivity from inferior frontal to medial temporal regions (IFC-MTC) to facilitate semantic access, suggesting a compensatory mechanism to provide the neural resources needed for LP performance. Similarly, Krieger-Redwood et al. (2019) highlighted that increased within-DMN connectivity between the right anterior temporal lobe (ATL) and medial prefrontal cortex (mPFC) might translate to inefficient semantic retrieval, reflecting older adults’ reduced ability to access the semantic store in a goal-directed manner.

At a structural or neuroanatomical level, our understanding of the white matter integrity that accompanies these midlife functional changes remains largely unexplored, despite recent research highlighting white matter degradation as an explanation for the onset of LP deficits (Kljajevic & Erramuzpe, 2019; Sánchez et al., 2023; Troutman et al., 2022; Yeske et al., 2021). Within the language network (Shekari & Nozari, 2023), microstructural integrity assessed via diffusion tensor-based metrics (e.g., fractional anisotropy, radial diffusivity) is associated with higher-order aspects of naming, such as lexico-semantic selection, notably along the left superior fasciculus (SLF III; Troutman & Diaz, 2020) and inferior longitudinal fasciculus (ILF; Kantarci et al., 2011; Stamatakis et al., 2011; Troutman & Diaz, 2020). Moreover, the frontal aslant tract (FAT; Rizio & Diaz, 2016; Troutman & Diaz, 2020), which connects the IFG to the premotor cortex (SMA and pre-SMA), is relevant for the sensorimotor aspects of fluency and speech-to-motor planning, with its anterior terminations associated with working memory (Varriano et al., 2020). This is in line with a recent study suggesting that converging locations between the left frontal aslant tract (FAT), frontostriatal tract (ST-FO; Stamatakis et al., 2011), and left SLF I are essential for efficient verbal working memory (Ribeiro et al., 2023). In addition, the middle longitudinal fasciculus (MLF) can also support the executive functions and working memory aspects of LP (Rizio & Diaz, 2016; Troutman & Diaz, 2020). Correspondingly, these studies align with recent findings (Roger, Rodrigues De Almeida, et al., 2022) on the left arcuate fascicle (AF), SLF III, ILF, and thalamic-premotor projections, which overlap spatially with language networks. In addition, spatial concordance was found between the left SLF II/left cingulum bundle (CG) and control-executive subnetwork, as well as between the fornix/hippocampal formation and the abstract-knowledge (DMN-related) subnetwork (see in detail Roger et al., 2022).

Based on behavioral and neurofunctional results, a recent study reported a comprehensive overview of various compensatory mechanisms for LP with aging (LARA model, Lexical Access and Retrieval in Aging, Baciu et al., 2021). This model describes two dimensions, LA (Lifespan Aging, uniform aging; natural effect of age) and RA (Reserve Aging, idiosyncratic aging; effect of cognitive reserve), according to two dimensions: Language-Specific (LS, specific to LA dimension) and Domain-General (DG, specific to RA dimension). Despite being grounded in experimental findings, the LARA model remains predominantly descriptive, and the brain structure dynamics that fit these dimensions are not included.

In the current study, our primary objective is to evaluate white matter changes supporting LP decline that emerges in midlife (Fargier & Laganaro, 2023; Oosterhuis et al., 2023). We anticipate decreased white matter integrity across the brain, especially along the dorsal and frontostriatal pathways, correlating with neurocognitive changes linked to age-related LP decline. Furthermore, we expect nonlinear microstructural changes that reflect accelerated cognitive decline in older adults. In line with the LARA model (Baciu et al., 2021), we further hypothesize that these white matter changes will align with two dimensions: (1) Lifespan Aging/Language-Specific LA/LS, involving semantic-related mechanisms with enhanced diffusivity along lateral pathways to enhance lexico-semantic representations, and along ventral mesial pathways to strengthen lexico-semantic memory retrieval. In line with Hoyau et al. (2018), this dimension could represent a compensatory mechanism that onsets in midlife to maintain LP performance, and (2) Reserve Aging/Domain-General RA/DG, representing the decline of domain-general mechanisms associated with reduced diffusivity within prefrontal regions, potentially involving SLF II and CG bundles. In line with Krieger-Redwood et al. (2019), this could challenge efficient and goal-directed access to the semantic store, compromising LP performance in older adults.

## 2. Material and Methods

Using state-of-the-art tractography techniques, we analyzed diffusion-weighted imaging and lexical production (LP)-related neuropsychological data of 155 healthy middle-aged adults (aged 45-60) from the CamCAN cohort (Cam-CAN et al., 2014). Figure 1 outlines the study workflow. After preprocessing diffusion-weighted imaging (DWI) data, we investigated the relationship between middle age and structural integrity by generating a track-weighted fractional anisotropy image (TW-FA) for each subject (Calamante, 2017). Subsequently, using Non-Negative Matrix Factorization (NMF), we reduced the dimensionality of these TW-FA images to identify sets of networks consisting of structurally covarying voxels. We then assessed the impact of age on the average microstructural integrity of each NMF network by using generalized additive models. Finally, we applied Partial Least Squares (PLS) correlation analysis at the voxel and network levels to examine how localized and distributed microstructural integrity changes covary with LP performance during middle age. To enhance the theoretical relevance of our study, a preliminary lifespan analysis (not depicted in Figure 1)) was conducted to specify the most critical age window for cognitive aging. This age window guided the main statistical analyses described earlier (please refer to section 2.3.1).

**Figure 1:**
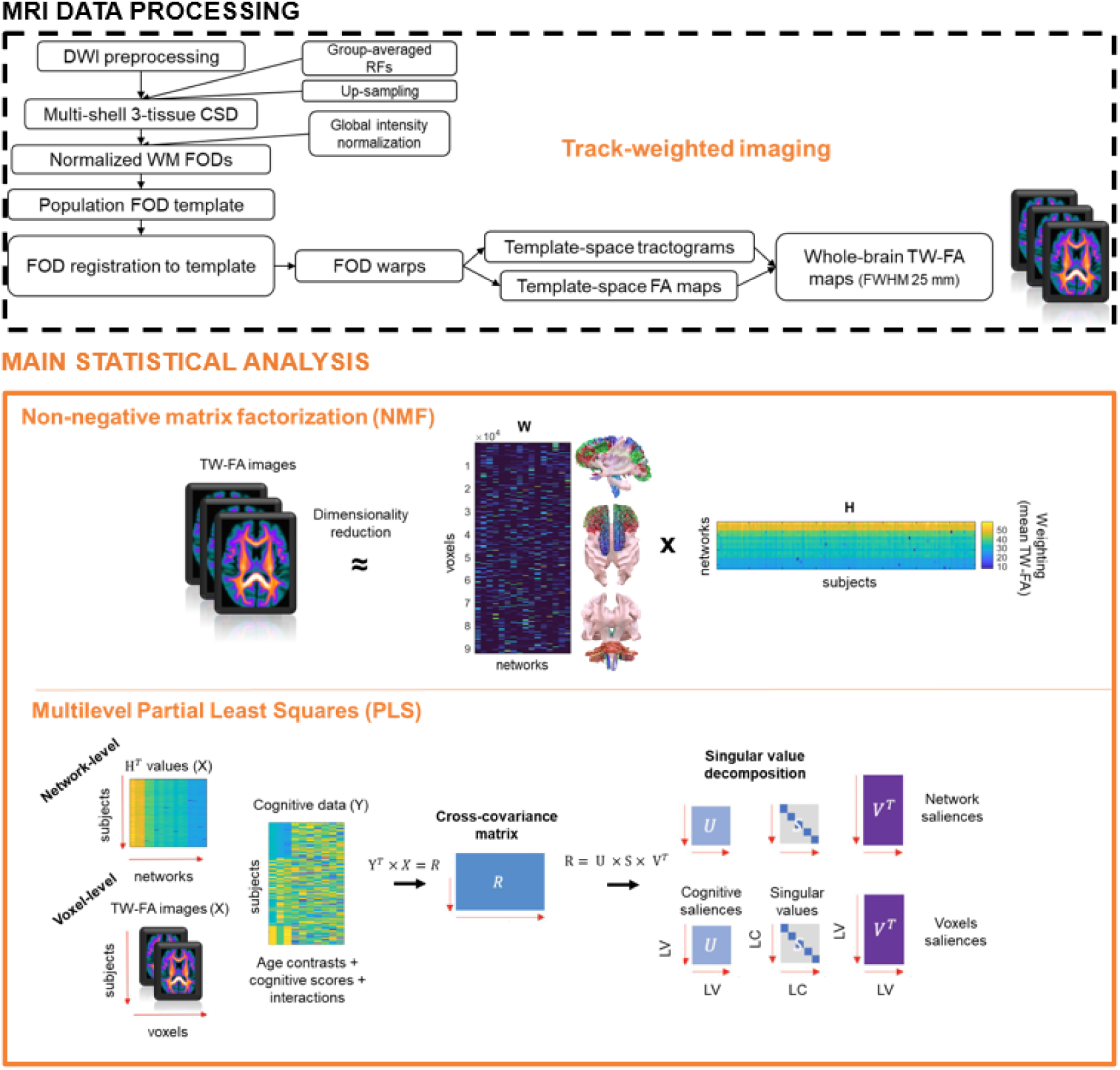
Workflow of the study. For multilevel PLS analysis, only one covariance matrix (R) is depicted for visual clarity; however, it is important to emphasize that a separate PLS model was executed using either voxel-level TW-FA values or network-level values derived from NMF (H values), resulting in two distinct covariance matrices.

### 2.1. Data and MRI acquisition

#### 2.1.1. Participants

We selected 155 healthy adults (76 females and 79 males) aged–45-60 at the time of MRI acquisition based on our preliminary analysis (refer to section 2.3.1). These individuals were chosen from the Cambridge Center for Ageing and Neuroscience project dataset (Cam-CAN et al., 2014). The East of England Cambridge Central Research Ethics Committee approved the CamCAN study. Please refer to Taylor et al. (2017) for further recruitment information. There was no significant difference in mean age at the time of MRI acquisition (*p*_FDR_ = .52) or handedness (*p*_FDR_ = .49), but middle-aged females tend to have better overall cognition (MMSE; t = 2.7, *p*_FDR_ = .017) across genders, and middle-aged males tend to have a higher total intracranial volume (t = -9.9, *p*_FDR_ < .001) (Figure A.1, Appendix A).

#### 2.1.2. Cognitive data

To assess lexical production (LP)-related performance, we selected eight neuropsychological tests that showed either: (i) a direct relationship with lexical production (LP) (e.g., Verbal Fluency, Naming, and Tip-of-the-Tongue tests) or (ii) an indirect relationship via the domain-general (DG) and language-specific (LS) mechanisms posited by the LARA model (Baciu et al., 2021). Table 1 shows the main cognitive processes associated with each test. Middle-aged females show better performances in Naming (t = 2.8, *p*_FDR_ = .04) and Story recall tasks (t = 2.3, *p*_FDR_ = .05). Specifically, the gap in sentence comprehension performances in favor of women grows as age increases (t = 2.18; *p* = .03) (see Table A.2, Appendix A for the results).

**Table 1.**
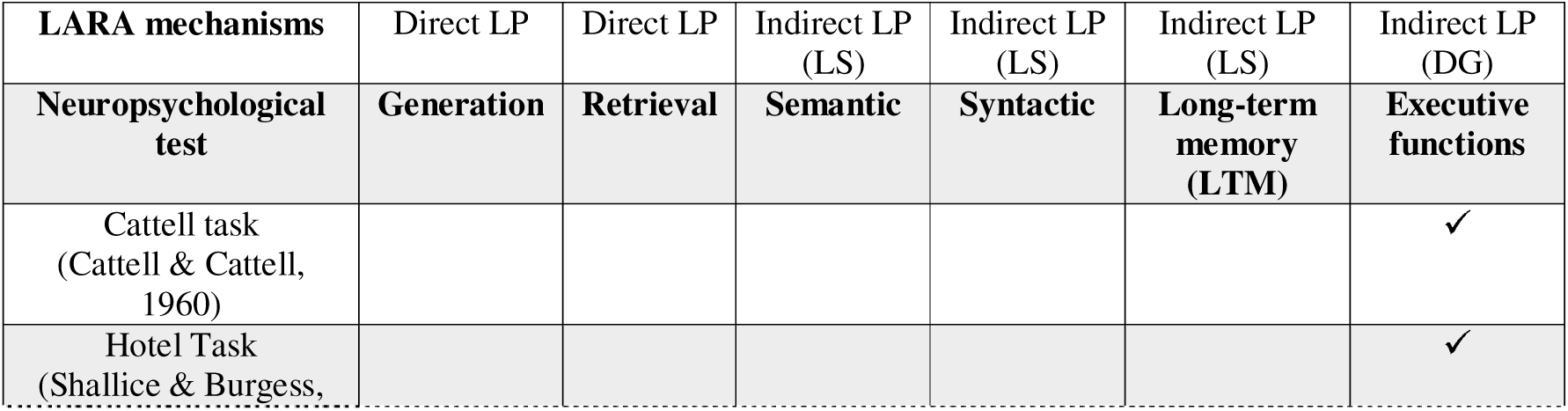
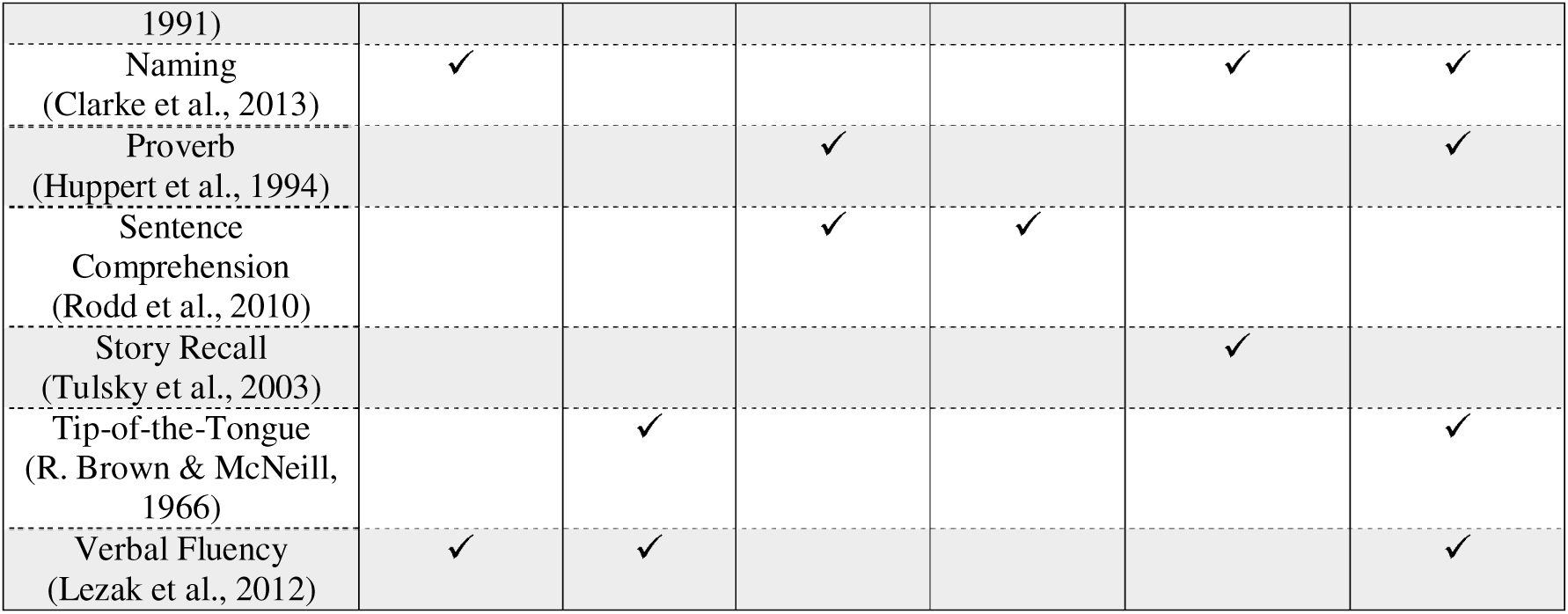
Presentation of the eight neuropsychological tests considered in this study and associated cognitive processes. A detailed description can be found in Table A.1, Appendix A, and the effect of age on cognitive performances is illustrated in Figure A.2, Appendix A.

#### 2.1.3. Diffusion MRI

Multi-shell diffusion-weighted imaging (DWI) data were acquired using a twice-refocused spin echo sequence. The acquisition parameters were as follows: 66 axial slices, TR/TE = 9100/104 ms, 2 mm isotropic voxels, 30 directions (*b*-values: 1000 and 2000 s/mm2), and three b0 images with an acquisition time of ∼10 min. Details can be found in the study by Taylor et al. (2017). All DWI data were preprocessed using MRtrix3 (Tournier et al., 2019) (version 3.0.4; https://www.mrtrix.org/).

### 2.2. MRI data processing

#### 2.2.1. Diffusion MRI

Preprocessing includes denoising (Veraart et al., 2016), Gibbs artifact removal (Tustison et al., 2010), eddy current and motion correction with FSL (Andersson et al., 2003), and bias correction with ANTs N4. DWI images were then upsampled to a voxel size of 1.5 mm^3^ using cubic B-spline interpolation (Raffelt et al., 2012). Following these preprocessing steps, fiber orientation distributions (FOD) were computed using the multi-shell 3-tissue Constrained Spherical Deconvolution model (MSMT CSD; Dhollander et al., 2016; Jeurissen et al., 2014; Tournier et al., 2007) with group-averaged response functions calculated before upsampling. Finally, a joint bias field correction and intensity normalization were applied to the resulting FOD images.

#### 2.2.2. Track-Weighted Imaging

We quantified structural integrity at each voxel location using Track-Weighted Imaging (TWI). Compared to conventional tensor-based metric, TWI enhances the sensitivity to microstructural changes and the representation of crossing fiber configurations (Calamante, 2017; Calamante et al., 2012). The TW images were generated in four steps. (1) Whole-brain tractography (iFOD2; 10 million streamlines, max. length = 250 mm, cutoff = 0.06, backtrack option) to obtain subject-specific tractograms while reducing reconstruction biases (SIFT2; Smith et al., 2015); (2) Generation of a study FOD template to ensure compatibility of tracks’ length and spatial location between participants (Willats et al., 2014). The template was based on normalized WM FOD images (Raffelt et al., 2011) from a subset of 35 participants, controlling for age (mean and standard deviation within 1%) and gender. Spatial correspondence across participants was achieved through FOD-guided nonlinear registration (Raffelt et al., 2011, 2012), producing a transformation image used to warp the tractograms to template space; (3) generation of subject-specific Fractional Anisotropy (FA) images using iterative least-squares (Basser et al., 1994; Veraart et al., 2013), and registered to the template space with the FOD-computed transformation images; (4) Generation of Track-Weighted FA images (TW-FA) by counting the number of tracks traversing a given voxel and weighting this count by the integrity (FA) of each track. We used a Gaussian neighborhood weighting of 25 mm (FWHM) to assess the short-to-mid-range variations in track structural integrity (Willats et al., 2014).

### 2.3. Statistical analysis

Statistical analysis was conducted in MATLAB (R2020b) and R (4.2.1). We controlled for handedness, gender, MMSE scores, and total intracranial volume in all the models (Eikenes et al., 2023).

#### 2.3.1. Preliminary analysis

To increase the theoretical relevance of our study, we determined the most relevant age window for cognitive aging with the hypothesis that middle age is a critical period.

*Method*. We modeled the trajectory of each cognitive score across the lifespan (18-88 years; 628 participants) using generalized additive models (GAM; Wood, 2006, 2017). We then calculated the curvature of each trajectory at each age point, selected the significant values (i.e., confidence intervals do not contain 0), and averaged them across the seven cognitive scores. The inflection point of cognitive aging corresponds to the point at which the curvature crosses zero (i.e., changes its sign). Computational details are available in supplementary material (Appendix A). *Results.* Age 53-54 is the average inflection point for LP-related performance This confirms that middle age is a critical period for cognitive aging and indicates a sudden acceleration in cognitive decline around this age, specifically driven by a drop in executive function performances (i.e., fluid intelligence via the Cattell task). After visually inspecting Figure A.3 (please refer to Appendix A), we selected a sample of middle-aged adults between the ages of 45 and 60 to target the period of transition associated with the onset of LP-related decline. Specifically, we chose at interval of 15 years that could equally be divided into 3 age groups for later statistical analysis (more details in section 2.3.3).

#### 2.3.2. Neuroanatomical level : Dimensionality Reduction

We explored the relationship between middle age and microstructural alterations in two steps. First, dimensionality reduction was used to identify a low-dimensional number of white matter networks. We then examined the effects of age on each network’s average microstructural integrity. All analyses were performed within a group mask to mitigate the spatial bias due to inter-individual variability and potential shortcomings of registration. Voxels with null TW-FA values across more than 5% of the participants were not included in the mask.

##### NMF network segmentation

We employed non-negative matrix factorization with an orthogonality constraint (https://github.com/asotiras/brainparts) to segment whole-brain white matter into networks. NMF is a dimensionality reduction technique widely used in neuroimaging to decompose high-dimensional datasets (e.g., voxel-level data) into meaningful networks (Sotiras et al., 2015). The orthogonality constraint ensured that networks were spatially non-overlapping. That is, each network comprised of a unique set of structurally covarying voxels, indicating that the microstructural integrity changes (TW-FA) along these voxels covary across participants (i.e., either strongly correlated or anti-correlated). Given its recent application in investigating age-related (Bagautdinova et al., 2023; Lynch et al., 2023; Sotiras et al., 2015) and biological processes captured by microstructural changes (Patel et al., 2023; Robert et al., 2022), and given that TW-FA values are non-negative, we considered NMF as an appropriate approach for our research question.

##### NMF implementation

NMF decomposes the matrix containing all TW-FA images (*A*) into two matrices: voxel matrix *W* (*voxels* × *k*) and subject matrix *H* (*k* × *participants*). The optimal number of networks *k* was determined using the procedure described by Bagautdinova et al. (2023): We performed NMF 10 times, incrementing the number of networks at each iteration (i) from *k=2* two *k=20* by steps of 2. The reconstruction error (*RE*), calculated as the Frobenius norm of the difference between the input and reconstructed matrix (Eq.2), guides the selection of the optimal *k*. The gradient of this error (*GradientRE*), representing the difference between reconstruction errors of two successive iterations (Eq.3), helped identify the point after which decomposing *A* into more networks yielded diminishing returns on model fit (Figure A.4, Appendix A).

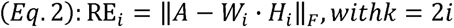

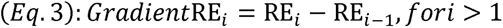

##### Network interpretation and composition

To accurately identify the brain pathways involved in each network, we automatically segmented 72 major bundles from the study template SH peaks using TractSeg (Wasserthal et al., 2018) and combined two metrics based on volumetric information (i.e., network and bundle-focused spatial overlap) with one metric based on fiber-connectivity information (Bullock et al., 2022). We considered brain pathways with a compositive score above or close to 1 as having a significant contribution. Further details are provided in supplementary material (Appendix A).

##### Age trajectory with GAM

To examine the effect of age, we fitted one GAM model for each network, with subject-level weightings (*H* values) as the dependent variable. These weightings reflect the average TW-FA values of each subject within a specific network. GAMs were fitted using restricted maximum likelihood (REML) for an unbiased estimate of fixed effects, a 3-knot spline to address overfitting concerns, and FDR-corrected (*q < 0.05)*.

#### 2.3.3. Neurocognitive level: Partial Least Squares

To examine the many-to-many relationships between age-related white matter integrity changes and LP performance, we employed a Partial Least Squares (PLS) correlation analysis using the toolbox myPLS (https://github.com/MIPLabCH/myPLS). PLS is a data-driven multivariate statistical technique that seeks to optimize the covariance between two datasets (Krishnan et al. 2011). In this study, PLS generates latent components that describe age-related patterns of covariance (i.e., coordinated changes with age) between Track-Weighted FA (TW-FA) values and cognitive scores related to lexical production (LP).

##### Multilevel PLS

While previous studies have used network-level NMF loadings in the PLS model (e.g., Patel et al., 2023; Robert et al., 2022), we reasoned that this may compromise the identification of more localized age effects at the voxel-level that would not have been captured using NMF as they may only appear when jointly considering white matter and cognitive performance changes. For this reason, we ran two models: network-level and voxel-level PLS. This multilevel approach allowed us to examine whether distributed (network-level) and localized (voxel-level) alterations are differently related to LP performance in middle age.

##### PLS setup

We organized the data into a structural (X) and cognitive (Y) matrix to set up the PLS. For X, TW-FA loadings (for voxel-level PLS) or NMF loadings (for network-level PLS; H values) were vectorized, stacked, and z-scored across participants so that each row encoded the values of one subject. For Y, cognitive scores were first quantile-normalized to improve Gaussianity (Pur et al., 2022) and then z-scored before prepending two age-related contrast variables and their respective interaction with the cognitive variables. The contrasts were set between three age groups: (contrast #1) 56-60 > 45-50 + 51-55, (contrast #2) 51-55 > 45-50. These contrasts ensured that nonlinear relationships with age were accounted for while remaining consistent with the parameter chosen previously for GAM modeling (i.e., 3-knot spline; see section 2.4.1). In this way, PLS allowed us to disentangle the common and differing covariance effects across age groups without disregarding absolute differences in either dataset (Zöller et al., 2017).

##### PLS implementation and interpretation

To start the PLS, the structural matrix (X) was cross-correlated with the cognitive matrix (Y), resulting in a covariance matrix (*R*). Singular value decomposition (SVD) was applied to decompose R, yielding a set of singular value (*S*), structural (*U*), and cognitive (*V*) saliences. The significance of *S* was assessed using 10,000 permutations and the robustness of salience (*U* and *V*) with 1000 bootstrap resamples. *S* contains singular values that represent the amount of information shared between the two datasets for each latent component (LC). *U* and *V* contain salience weights representing the contribution of each voxel or cognitive variable to the pattern of covariance captured by each LC. For each LC, we obtained two latent variables, one structural (LV_struct_) and one cognitive (LV_cog_), by projecting the saliences back onto the corresponding z-scored dataset (i.e., LV_struct_ *= U*·X and LV_cog_ *= V^T^*·Y). To assess the robustness of structure and cognitive saliences, we report bootstrap sampling ratios (BSR), calculated as the salience weight over its bootstrapped standard deviation. A high BSR (± 2.58) indicates a robust contribution within a 99% confidence interval (Krishnan et al., 2011).

## 3. Results

### 3.1. Neuroanatomical level

We applied non-negative matrix factorization (NMF) to identify patterns of white matter microstructural integrity (WMI) changes in middle-aged adults. We identified a total of 16 white matter networks. Six white matter networks showed significant modifications with age, indicating a broad structural reorganization between 45 and 60.

The average WMI of 4 of these 6 networks decreased nonlinearly with age, confirming that middle age is a transition period at the structural level. Specifically, we observed the largest decrease in WMI beyond age 53, compromising widely distributed brain pathways. Figure 2 and the following white matter network presentation are organized based on the extent of change with age, ranging from the most to least significant.

**Figure 2:**
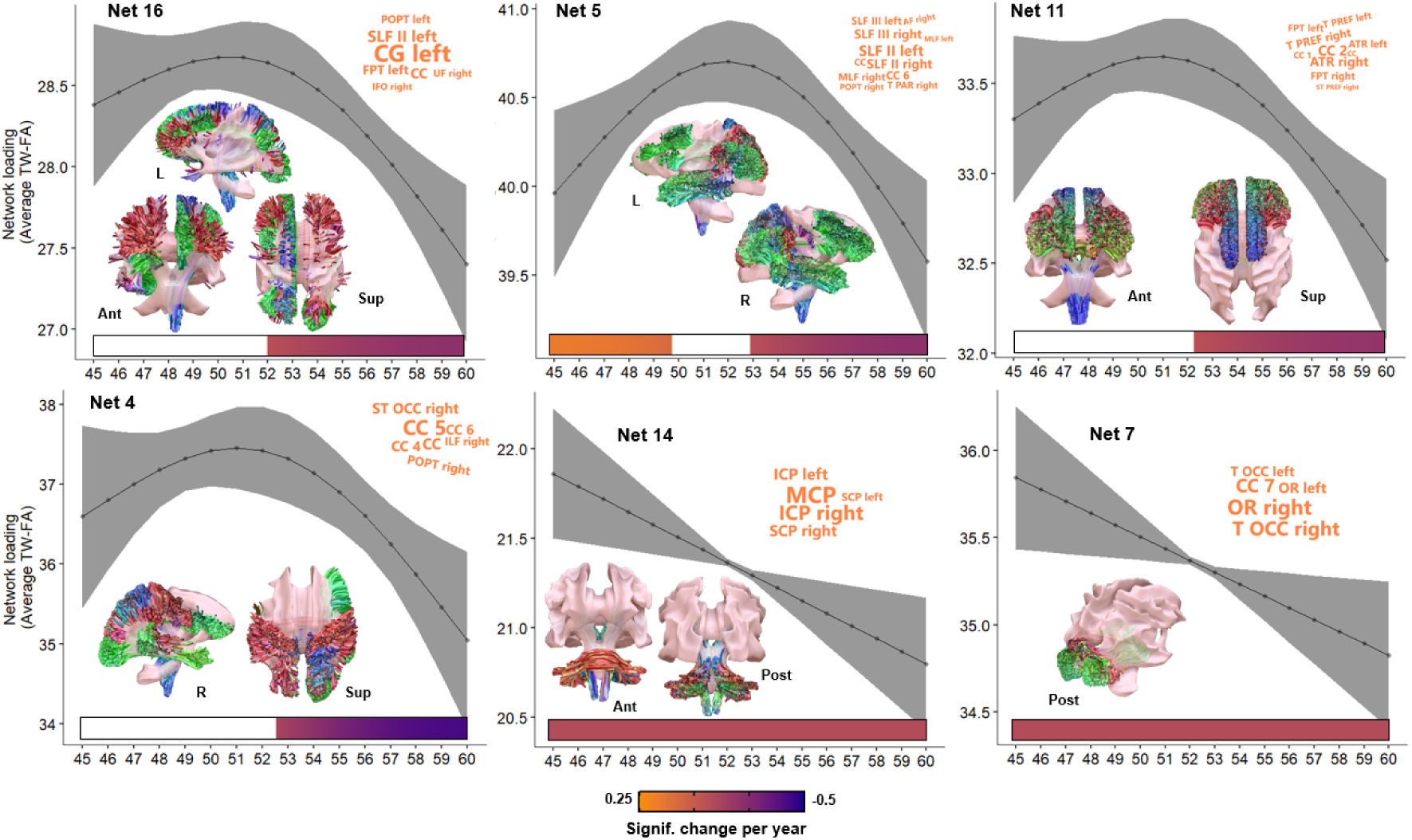
Age-related white matter networks. **Age trajectory:** Plots display the significant relationships between TW-FA (track-weighted microstructural integrity) and age quantified by generalized additive models. Bars above the x-axis represent the derivative of the fitted GAM smooth function and correspond to age windows of significant changes. The color of this bar indicates the magnitude of the TW-FA change with age: orange when increasing (i.e., positive derivative), purple when decreasing (i.e., negative derivative), and white when no significant change was found. A derivative was considered significant when its confidence interval did not contain zero. The networks are sorted from the one showing the strongest (top left) to the one showing the weakest age-related effect (bottom right). **Word cloud:** Depicts the major brain pathways a given network engages (see Appendix A for details on calculations). Tract names correspond to the TractSeg nomenclature (https://github.com/MIC-DKFZ/TractSeg). **Brain illustration.** Depicts the bundle fibers (min. length 25 mm) crossing one or many of the voxel clusters associated with a given network. Fiber orientations: red (left-right), green (anterior-posterior), and blue (superior-inferior). Displayed on a mesh of the average TW-FA image using the Surf Ice software (https://www.nitrc.org/projects/surfice/). *Abbreviations:* L (left), R (right), Ant (anterior), Post (posterior), Sup (superior), Inf (inferior).

In the discussion section, we discuss extensively how the following Net 16, 5, and 11 may contribute to the cognitive control processes required for lexical production. Specifically, **Net 16** shows reduced WMI from age 52 onwards (F = 7.6, *p*_FDR_ = .016, partial R² = .09), primarily in a distributed left fronto-parietal-pontine network. This impacts the left-hemispheric integrity of the left cingulum bundle (CG: z-scored contribution = 2.46), SLF II (1.61), left cortico-pontine tracts (FPT: 1.3; POPT: 1.13), also engaging the corpus callosum (CC: 1.42) and minor contributions from the right UF and right IFO (1.03/0.98). **Net 5** is a bilateral association network with slight asymmetry along a left-right dorso-ventral axis, showing an initial increase in WMI (age 45-50) followed by a decrease from age 53 onwards (F = 5.73, *p*_FDR_ = .016, partial R² = .08). Specifically, this translates to coordinated white matter modifications between the dorsal SLF II left/right (1.91/1.68) and the more ventral SLF III left/right (1.39/1.63) & MLF left/right (0.98/1.35). This network is bound across hemispheres via the isthmus of the corpus callosum (CC 6: 1.48). We also note minor contributions from the right thalamo-parietal (T PAR: 1.23), right POPT (1.12), and right AF (1.12). **Net 11** primarily shows reduced thalamo-(pre)-frontal WMI, with a slight dominance of the right-hemispheric tracts, beyond age 52 (F = 6.72, *p*_FDR_ = .016, partial R² = .08). This impacts the integrity of the anterior thalamic radiations (ATR; left: 1.48, right: 1.9), fronto-pontine pathways (FPT; left: 1.43, right: 1.49), thalamo-prefrontal (T PREF; left: 1.4, right: 1.49) and right striato-prefrontal (ST PREF; 0.97) bound mostly by the genu and rostrum of the corpus callosum (CC2: 2.06; CC1: 1.25).

We identified 3 additional networks that were only moderately associated with middle age: **Net 4** shows a moderate loss of WMI beyond age 52 as well (F = 5.08, *p*_FDR_ = .045, partial R² = .06) in a distributed right network. This primarily affects the communication of the anterior/posterior midbody of the CC (i.e., CC4/primary motor: 1.24, and CC5/primary somatosensory: 1.83), the isthmus (CC6: 1.19), the right striato-occipital (ST OCC; 1.25), right POPT (1.07), and right ILF/right AF (0.98/0.95). **Net 14** and **Net 7** were associated with steady loss of integrity respectively engaging inferior (left: 2.84, right: 3.69), middle (4.13), superior (2.01/2.77) cerebellar peduncles (F = 8.03, *p*_FDR_ = .02, partial R² = .05), and visual-occipital pathways via the optic radiations (left: 1.6, right: 2.3) and T OCC (1.49/2.19) bound together by the splenium of the corpus callosum (CC 7: 2.08) (F = 5.76, *p*_FDR_ = .048, partial R² = .04).

### 3.2. Neurocognitive level

To evaluate the white matter changes supporting the emergence of LP difficulties in middle age, we performed two partial least squares (PLS) analyses: one at the network level to examine the effect of distributed alterations (i.e., network-level PLS) and one at the voxel level to examine the effect of localized alterations (i.e., voxel-level PLS).

In line with our hypothesis, multilevel PLS found two latent components consistent with a domain-general (LC1) and language-specific mechanism (LC2) (Figure 3A). LC1 was significant at both levels of analysis (51.73/19.57% of the total shared variance; singular value = 228.4/9581.1; *p_Bonferroni_* = .001/*p_Bonferroni_* = .002), and LC2 remained significant only at the voxel level after Bonferroni correction (13.76%; 8033.3; *p_Bonferroni_* = .002). The subsections below describe the cognitive and microstructural changes of each component and how their joint dynamic may suggest a relationship between middle-aged LP and the control of semantic representations (interplay LC1-LC2).

**Figure 3:**
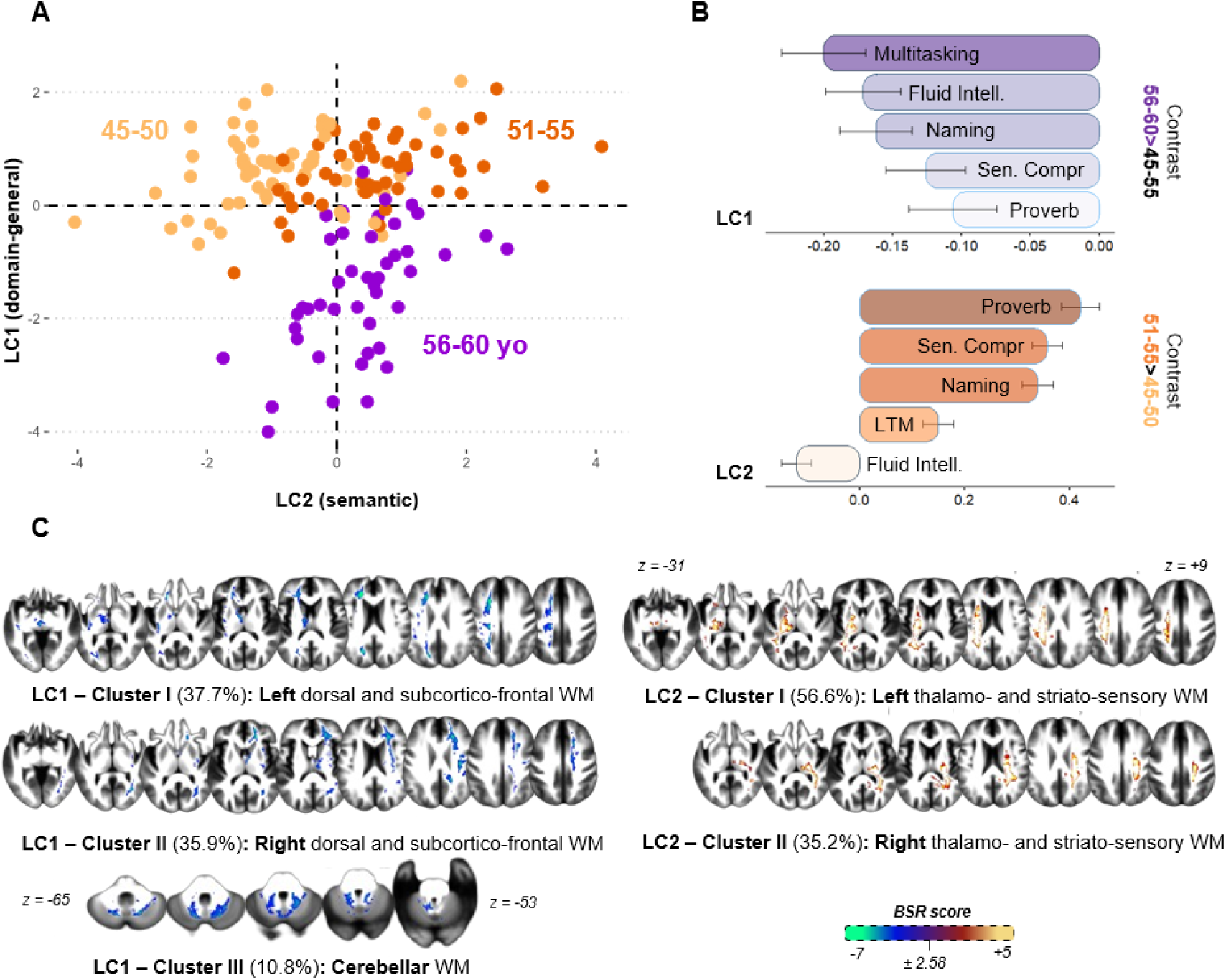
Results from voxel-level PLS analyses. **(A) Depicts the 2D latent space across middle-aged groups.** For each latent component (LC1 and LC2), latent cognitive and structural scores were combined with a PCA for visualization. See Figure B.3, Appendix B for the detail on each component and Figure B.1-2, Appendix B for diagnostic plots **(B) Cognitive saliences.** Only the variables with a robust contribution to the latent component are reported (i.e., bootstrap sampling ratio ± 2.58). Bars extend to the corresponding salience weight (*U*), and error bars indicate the bootstrapped standard deviation. **(C) Brain saliences.** Blue/green indicates a decrease in white matter microstructural integrity, and red/yellow indicates an increase. Please see Figure B.4, Appendix B for the raw brain saliences. Displayed on the study template is the average TW-FA image. Cluster analysis was performed using the MATLAB *bwconncomp* function. Voxels were considered part of the same cluster if they were both salient and connected in one of these directions (in/out, left/right, up/down; 3D connectivity set to 6). *Abbreviations*: bootstrap sampling ratio (BSR), white matter (WM).

#### 3.2.1. Latent component 1: Domain-general mechanism

The first latent component (LC1) aligned with our previous finding, suggesting that the magnitude of distributed and localized microstructural integrity alterations increases significantly in late middle-aged individuals (BSR_56-60>45-55_ = -9.66 for network-PLS/-19.2 for voxel-PLS) and correlated with an acceleration in cognitive decline (correlation between LVs: .42/.55).

##### Voxel-level PLS

At the cognitive level, individuals aged 56-60 versus younger adults (contrast 56-60>45-55), had lower performances in tasks related to executive functioning (see Section 2.1.2): multitasking (BSR = -6.59), fluid intelligence (-6.28), directly correlating with direct LP difficulties (naming = -6.22). More marginally, we also found a decline in sentence comprehension (-4.34) and proverb comprehension (-3.32). In comparison, individuals aged 51-55 versus younger middle-aged adults (contrast 51-55>45-50) exhibited a tendency for long-term memory (-2.73) and verbal fluency difficulties (2.69) (Figure 3B).

At the structural level, late middle-aged individuals showed a significant bilateral reduction in microstructural integrity in three localized clusters of white matter accounting for 84.39% of all alterations: Cluster I (37.65%) and Cluster II (35.91%) primarily engaged either left or right, subcortico-frontal and dorsal pathways; Cluster III engaged the cerebellar white matter (10.82%) (see Table 2 & Figure 3C).

**Table 2.**
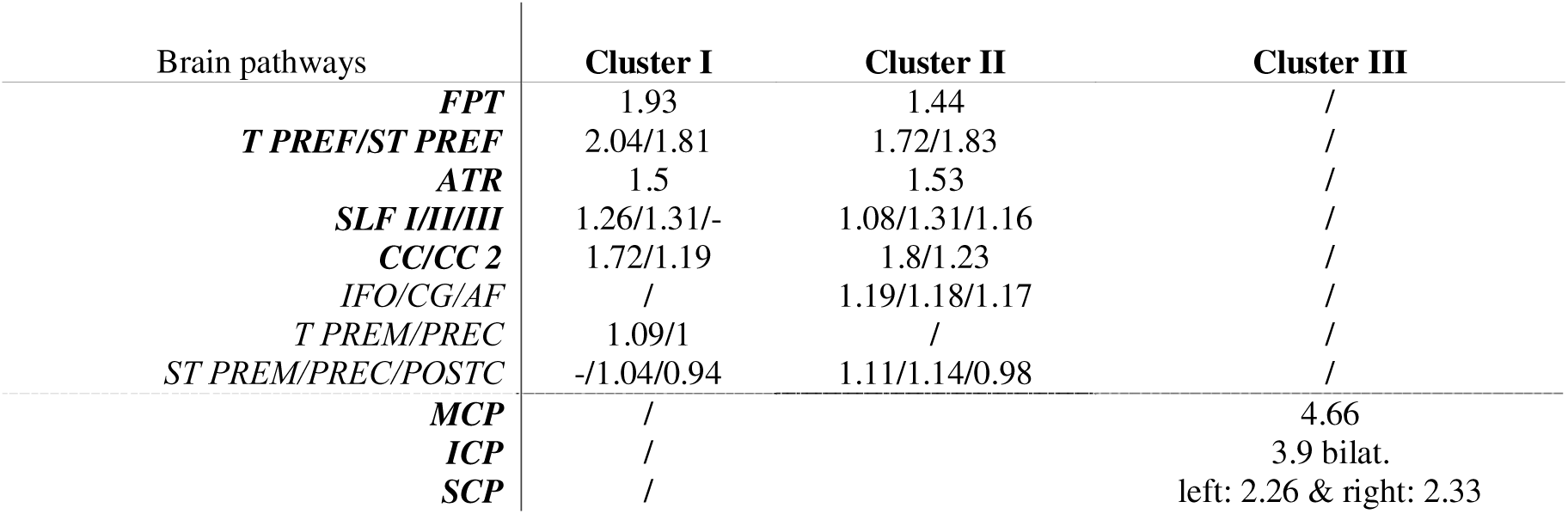
Brain pathways contributing to LC1. Numerical values are a z-scored composite metric based on volumetric and connectivity-based overlap with the robust brain saliences (please refer to Appendix A for details). A score above 1 shows a significant contribution. Pathways highlighted in bold font show the highest contribution. Tract names correspond to the TractSeg nomenclature (https://github.com/MIC-DKFZ/TractSeg).

##### Network-level PLS

In line with these results, we found corresponding distributed alterations in all 6 age-related networks (Net 16: -7.21; Net 11: -6.25; Net 5: - 6.18; Net 4: -3.73; Net 7: -3.55; Net 14: - 3.15; Figure 4B). Specifically, Net 16 (left fronto-parieto-pontine), 11 (thalamo-frontal) and 7 (visual-occipital) robustly underpinned the onset of multitasking difficulties in late middle-age (BSR = -3.2). Net 5 (bilateral association) additionally covaried with verbal fluency abilities and modulation of Net 14 (cerebellar) seemed to underpin a wider range of LP-related tasks except for multitasking (Figure 4A).

**Figure 4:**
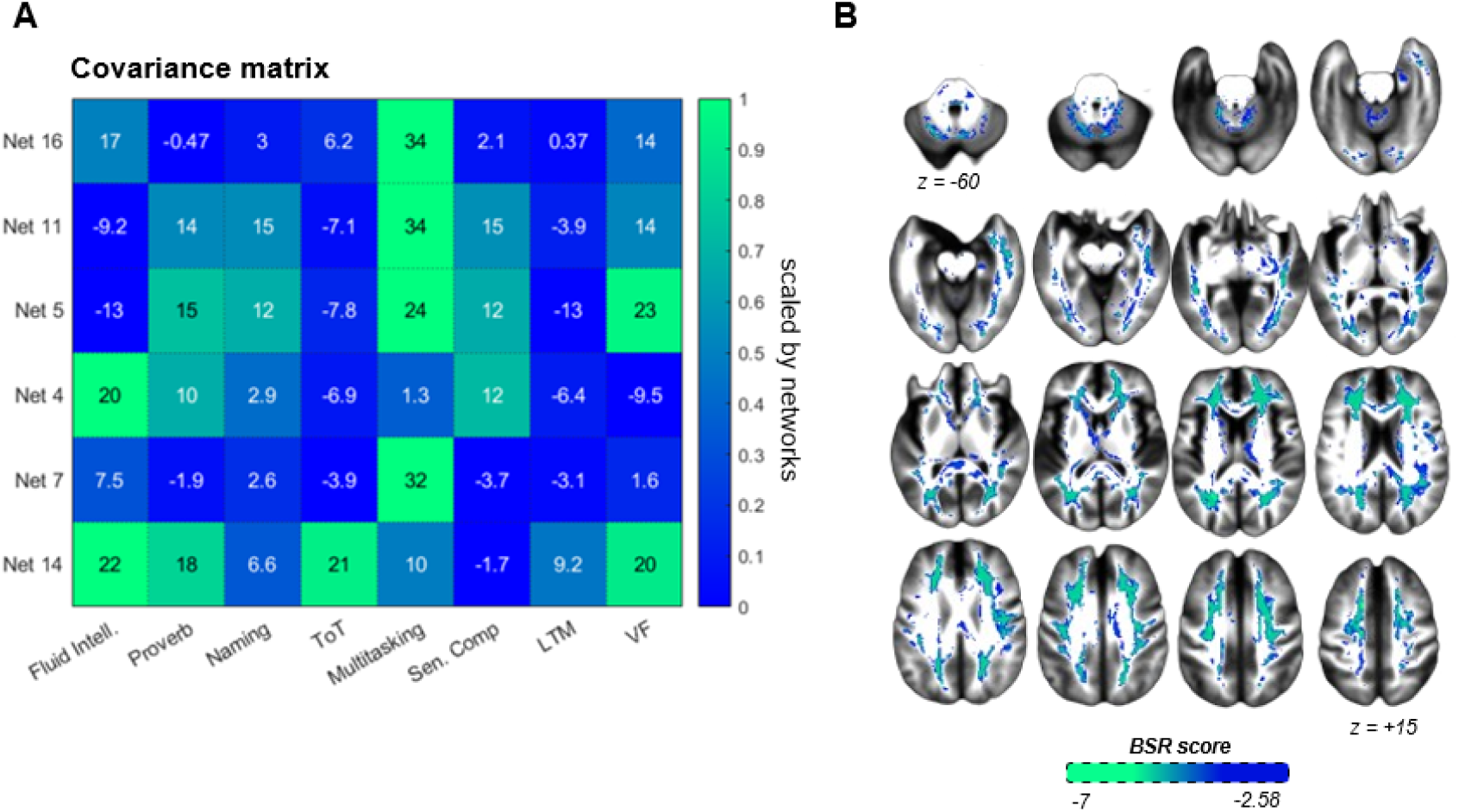
Results from network-level PLS. (A) Covariance matrix. (B) Brain structural saliences. Displayed on the study template is the average TW-FA image. *Abbreviations*: bootstrap sampling ratio (BSR).

Moreover, we found a robust contribution from networks not previously associated with age (primarily Net 8, Net 9, and Net 15 covarying with multitasking and long-term memory; see Figure B.5, Appendix B). This is in line with high residual variability found along the latent structural variable (x-axis), especially among late middle-aged adults (Figure B.3, Appendix B), suggesting that (unmodeled) age-invariant factors could primarily affect the microstructural integrity of white matter networks.

#### 3.2.2. Latent component 2: Semantic mechanism

The second latent component (LC2) differed markedly from LC1, revealing an intriguing age-related trajectory stabilizing around aged 51-55 (BSR_51-55>45-50_ = 12.41).

##### Voxel-level PLS

At the cognitive level, individuals aged 51-55 compared to early middle-aged adults showed sign of increased semantic knowledge (proverb = 14.76; sentence comprehension = 12.41), translating to better naming (10.02) and long-term memory performances (4.99), and this increase was proportional to the decline in fluid intelligence (-3.88). Interestingly, despite similar activation of semantic representations, late middle-aged adults exhibited significant multitasking (-6.7), naming (-4.56), and semantic abstraction difficulties (-4.18), suggesting that the onset of cognitive control challenges reported in LC1 could regulate the access to semantic representations (Figure 3B).

At the structural level, this pattern of cognitive changes covaried with an increase in microstructural integrity in two clusters of white matter accounting for 81.73% of all modifications: Cluster I (56.5%) and Cluster II (35.23%) primarily engage either left or right thalamo- and striato-sensory pathways (see Table 3 & Figure 3C).

**Table 3.** Brain pathways most contributing to LC2. Numerical values are a z-scored composite metric based on volumetric and connectivity-based overlap with the robust brain saliences (please refer to Appendix A for details). A score above 1 shows a significant contribution. Pathways highlighted in bold font show the highest contribution. Tract names correspond to the TractSeg nomenclature (https://github.com/MIC-DKFZ/TractSeg).

| Brain pathways | Cluster I | Cluster II |
| --- | --- | --- |
| **STR** | 2.22 | 2.54 |
| **T PREC/ST PREC** | 2.05/1.78 | 2.09/1.71 |
| **T POSTC/ST POSTC** | 1.88/1.6 | 2.12/1.74 |
| **FPT** | 1.71 | / |
| **POPT** | 1.78 | 2 |
| **CST** | 1.82 | 1.62 |
| **T PREF/PREM** | 1.32/1.3 | / |
| **T PAR/ST PAR** | 1.54/1.28 | 2.11/1.88 |
| **SLF II** | 1.01 | 1.18 |

#### 3.2.3. Joint dynamic of LC1 and LC2: Cognitive control over semantic representations

Having established the two structure-cognition patterns of covariance in middle age, we sought to gain additional insight by considering their joint dynamic. Doing so points to an interesting dynamic characterized by divergent middle-aged trajectories: executive functions tend to decline but semantic knowledge accrues. Moreover, these trajectories intersect around age 53-54 (Figure 5), which concurs with the inflection point for accelerated LP-related decline and microstructural integrity damage respectively reported in the preliminary (Section 1.3.1) and neuroanatomical analysis section (Section 2.1).

**Figure 5.**
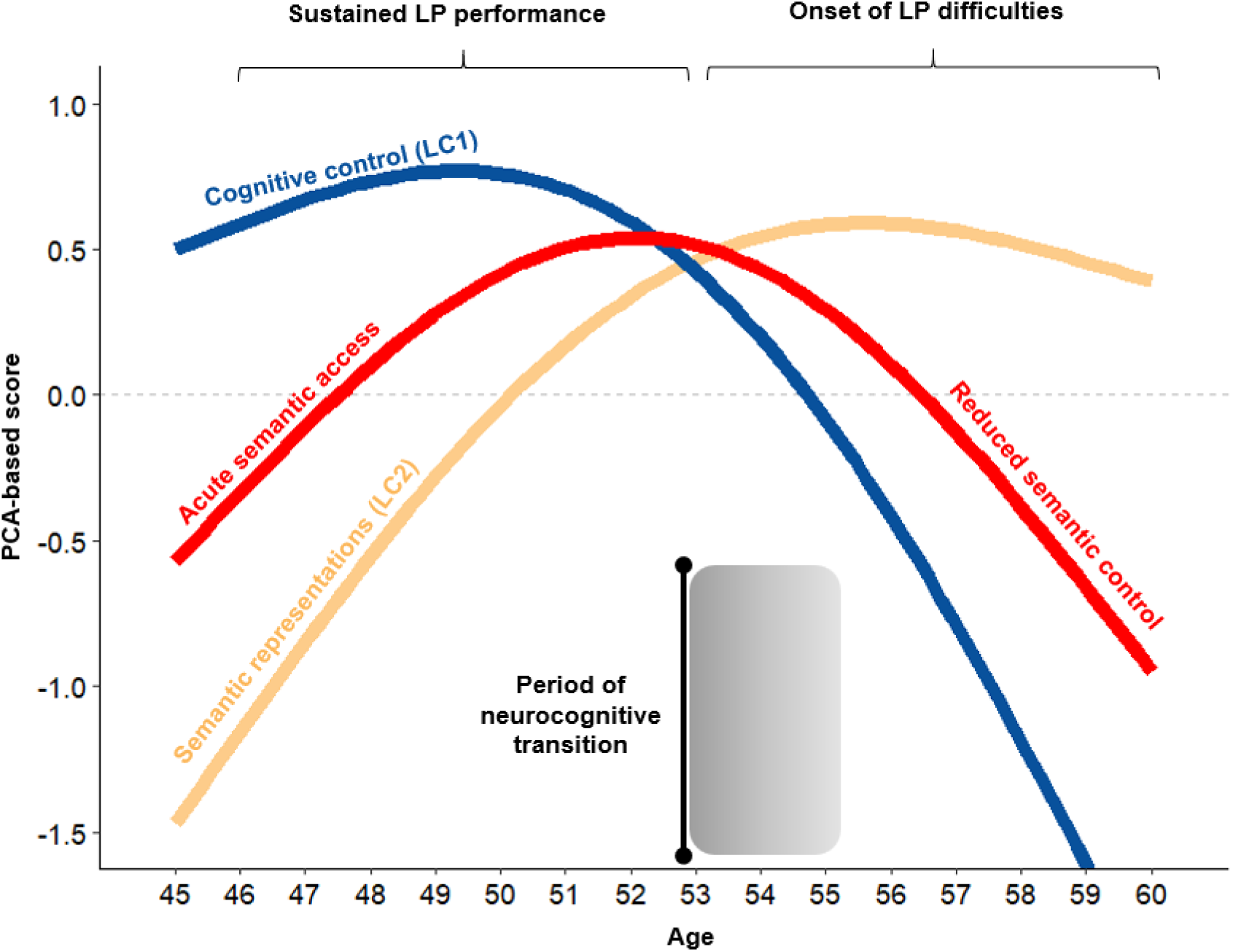
Compensatory dynamic of lexical production difficulties in midlife. Each component’s latest cognitive and structural variables were combined with a PCA for visualization. Bars above the x-axis represent the second-order derivative of the fitted GAM smooth function in red. A blue hue indicates a period of transition during middle age. A derivative was considered significant when its confidence interval did not contain zero.

## 4. Discussion

This study evaluated white matter changes supporting lexical production (LP) decline that emerges in middle age. Indeed, healthy aging is associated with a gradual decline in LP, beginning in middle age and manifesting in increased naming latencies. To delay the onset of LP decline as much as possible, middle-aged adults may rely on compensatory mechanisms, as proposed by the LARA model (Lexical Access and Retrieval in Aging; Baciu et al., 2021). However, our knowledge of the white matter changes supporting these mechanisms remains incomplete. Therefore, in this study, we employed advanced machine learning approaches on diffusion-weighted imaging and neuropsychological data from a sample of 155 middle-aged adults to (i) identify the brain pathways undergoing age-related microstructural changes and (ii) explore how LP performance is related to structural brain reorganization as individuals age.

Consequently, our main findings can be presented at two levels: (a) At a neuroanatomical level, middle age compromises the microstructural integrity of distinct white matter networks, revealing novel associations between brain pathways. (b) At a neurocognitive level, we reaffirm that middle age is a critical period for language processing. Indeed, while early middle-aged adults may rely on significant semantic access to maintain LP performance, our study suggests that late middle-aged adults may fail to exert cognitive control over these semantic representations, eventually leading to word-finding difficulties. The present study furthers our comprehension of middle-aged brain structure changes and sheds light on the neurocognitive transition leading to the onset of LP decline in middle age. We will discuss our neuroanatomical findings and the neurocognitive changes related to LP in middle-aged adults.

From a neuroanatomical standpoint, our results reveal that large-distributed WM pathways undergo coordinated microstructural changes with age. This suggests that short-to-mid-range (i.e., ∼25 mm length) microstructural alterations along known brain pathways can be grouped and analyzed within WM networks. Crucially, this network-level approach complements the conventional view that the age-related loss of integrity takes place along dorsoventral (Boban et al., 2022; Zahr et al., 2009) and antero-posterior gradients (Bennett et al., 2010; Boban et al., 2022). Indeed, our results reveal that the SLF (II/III) and thalamo-frontal pathways, often attributed with the largest age-related changes (Bonifazi et al., 2018; Hughes et al., 2012; Troutman et al., 2022), are part of heterogeneous WM networks bind together by callosal fibers: the posterior midbody (CC6) binds association pathways (SLF II and SLFIII/MLF; Net 5), and the rostrum and genu (CC1/2) bind anterior cortico-subcortical pathways (e.g., ATR, T PREF, FPT; Net 11). Temporally, the WM networks most affected by age (i.e., Net 16, 5, 11, 4) showed the largest microstructural alterations occurring from age 53 onwards. This aligns with previous studies concluding that discontinuities (i.e., nonlinear trajectory) in brain structure are likely to onset at middle age (Beck et al., 2021; Park & Festini, 2016) and further reinforces the idea that midlife is a critical period of major brain structural reorganization in line with reports on brain function and language performance (Deery et al., 2023; Hennessee et al., 2022; Park & Festini, 2016; Roger et al., 2023; Shafto & Tyler, 2014). Overall, our study supports the current evidence that dorso-ventral association (SLF II/III, MLF) and anterior cortico-subcortical pathways suffer the largest age-related microstructural alterations (Gunning-Dixon et al., 2009; Park & Festini, 2016; Yeatman et al., 2014). Specifically, the reduction of TW-FA in the anterior corpus callosum (CC1/2) and deep (pre-)frontal regions suggests that fibers of the forceps minor could be preferentially affected (Hsu et al., 2008; Kelley et al., 2019), along with critical age-related modulation of the integrity of fronto-limbic, and fronto-parieto-pontine fibers (i.e., FPT, POPT). Nonetheless, we also note that more posterior brain pathways do not show evidence for such disruptions. That is, cerebellar (Net 14) and visual-occipital (Net 7) pathways tend to exhibit a weaker and continuous (i.e., linear) loss in microstructural integrity as individuals age.

From a methodological standpoint, our results advocate for a network-level approach to complement anatomically defined bundle atlases (Bullock et al., 2019; Bullock et al., 2022; Catani & Thiebaut De Schotten, 2008; Mori et al., 2008; Rojkova et al., 2016; Wasserthal et al., 2018), leading to novel associations between brain pathways free from explicit spatial constraints (Bagautdinova et al., 2023). Ultimately, this emphasizes the need to apply advanced data-driven tools (i.e., dimensionality reduction via NMF; Sotiras et al., 2015) to capture the complexity of WM changes. This opens new research avenues for building complex yet highly interpretable models to explore how brain structure constrains brain function, for example projecting the functional signals onto white matter pathways of interest (see the functionnectome by Nozais et al., 2021, 2023) and, more broadly, assess the coupling between microstructure and brain function (e.g., Patel et al., 2023) (see also Calhoun & Sui, 2016 for a review on data fusion methods).

From a neurocognitive standpoint, our analyses offer new insights into the brain structure dynamics involved in maintaining lexical production (LP) performance in middle age. In line with our hypothesis, two latent components, LC1 and LC2, were identified, corresponding to the Domain-General (DG) & Language-Specific (LS) mechanisms posited in the LARA model (Lexical Access and Retrieval in Aging; Baciu et al., 2021).

On one hand, LC1 indicated a decline in DG processes in late middle-age, such as multitasking and fluid intelligence, coinciding with the onset of LP difficulties, particularly in naming. This aligns with evidence linking the speed of lexical retrieval and the generation of lexical predictions to fluid processing abilities (Brothers et al., 2017; Strijkers et al., 2011). Specifically, this suggests that LP demands substantial cognitive effort, as older adults’ reduced ability to flexibly allocate attentional resources (i.e., fluid processing; Carpenter et al., 1990) correlates with age-related LP decline. Structurally, LC1 highlighted microstructural integrity loss in bilateral medio-lateral WM areas, involving the association (SLF I/II/III) and anterior cortico-subcortical pathways (e.g., T PREF/ST PREF, ATR) that contribute to the white matter networks most affected by age (i.e., Net 16, 11, 5). Relatedly, a wealth of evidence concludes that such WM networks implement cognitive control mechanisms necessary for LP: (i) the SLF II and CG bundle along the anterior cingulate area (Net 16) (Hafkemeijer et al., 2014; Roger, Rodrigues De Almeida, et al., 2022), with the cingulum bundle being closely linked with inhibition tasks (Ribeiro et al., 2023); (ii) the anterior cortico-subcortical pathways (Net 11), especially right striato-prefrontal pathways (Buckner, 2004; Webb et al., 2020) and the right anterior thalamic radiations for response inhibition (Ribeiro et al., 2023); (iii) the SLF II and MLF for mental flexibility (Net 5) (Ribeiro et al., 2023; Rizio & Diaz, 2016; Troutman & Diaz, 2020). The link between DG and LP is further evidenced by a recent longitudinal study showing that the inability of structural brain networks to facilitate effortful brain state transitions is primarily associated with a deficit in executive functions (Tang et al., 2023). Of note, LC1 was also associated with minor sentence comprehension difficulties in late middle age. This could further underscore the high cognitive demands of LP, considering that lexical prediction is influenced by top-down comprehension strategies (Brothers et al., 2017).

On the other hand, LC2 revealed enhanced sentence comprehension, semantic abstraction, and long-term memory processes up until age 53-54. Interestingly, these changes were proportional to both (a) the increase in naming performances and (b) the decline in fluid abilities faced in early middle age. Given the constraint that fluid processing places on LP, this suggests that such an increase in semantic representations could provide the basis of a compensatory “semantic strategy” to maintain LP performance despite a decline in fluid-related abilities (e.g., problem-solving, learning). This is consistent with the idea that older adults may compensate for age-related decline in “control over LP” with higher vocabulary knowledge (Gollan & Goldrick, 2019). This also confirms that semantic knowledge accumulated over the lifespan (Salthouse, 2019; Spreng & Turner, 2019) is a viable alternative to meet reduced fluid processing with increased associative processing, thus lowering the control demands during LP (Spreng & Turner, 2021). Structurally, this tendency towards an increasingly “semanticized” cognition (Spreng et al., 2018) is associated with enhanced microstructural integrity in a left-dominant cortico-subcortical network (e.g., STR, T/ST PREC, T/ST POSTC), aligning with the idea that thalamocortical pathways are crucial for stable semantic representations at the semantic-lexical interface (Crosson, 2021).

Critically, our investigation sheds light on the interplay between DG and LS during middle age, suggesting that sufficient cognitive control is key to the compensatory “semantic strategy”. Indeed, in late middle age, LP decline co-occurs with significant DG deficits and increased semantic abstraction difficulties in LS. Yet, the brain structural changes of LS in late middle-aged adults remain comparable with that of younger adults (age 51-55), indicating late middle-aged difficulties are not underpinned by reduced semantic knowledge. Said differently, a richer repertoire of stored semantic resources in middle age, as captured by LS, appears to be a necessary but insufficient condition to maintain LP performance in late middle age. Instead, we argue that the compensatory dynamic of middle-aged LP is driven by the ability to use flexibly and abstract this semantic knowledge, that is the DG-LS interplay (as depicted in red in Figure 5). This supports the idea that semantic cognition depends on both representational and control neural systems (Hoffman & MacPherson, 2022; Wu & Hoffman, 2023), with semantic control regulating access to semantic representations (Branzi & Lambon Ralph, 2023; Ralph et al., 2017).

Taken together, our investigation provides empirical evidence in favor of the LARA model (Baciu et al., 2021) by suggesting that the interplay between domain-general (DG) and language-specific (LS) mechanism is crucial to understanding the onset of word-finding difficulties in the middle age. Importantly, this reaffirms that LP is a highly integrative cognitive function and the product of intra-(LS) and extra-(DG) linguistic processes (Hagoort, 2019; Hertrich et al., 2020; Roger, Banjac, et al., 2022). While middle-aged adults may adopt a compensatory “semantic strategy” to delay the onset of word-finding difficulties, this strategy may be compromised in late middle age when the ability to exert cognitive control over semantic representations is challenged, translating into a poorer filtering of irrelevant semantic associations (i.e., reduced semantic selection; Badre & Wagner, 2007; Barba et al., 2010; Jefferies, 2013).

Taking a broader perspective, this transition in midlife echoes recent proposals advocating for a learning-to-prediction shift (R. M. Brown et al., 2022) from an explorative to an exploitative cognitive mode (Hills et al., 2015; see Spreng & Turner, 2021 for a review), ultimately disproving that cognitive aging is solely synonymous with cognitive decline (Spreng & Turner, 2019). Indeed, exploiting accumulated semantic knowledge resources can be viewed as a predictive process, providing sufficient flexibility for comparing sensory inputs against long-term memory traces when given ambiguous, noisy, or incomplete lexical information (Bar, 2007; Bubic, 2010). In the context of LP, this signifies that middle-aged adults are better equipped to produce utterances elicited by semantic and/or episodic associations (i.e., inferential naming) compared to a typical picture naming task (i.e., referential naming) (Fargier & Laganaro, 2023). In the context of semantic processing, this is consistent with studies reporting that older adults make less use of contextual information as noted by Jongman & Federmeier (2022).

### Limitations and Perspectives

We mention that we did not explicitly model cognitive reserve factors despite their key role in capturing the inter-individual variability in language processing (Chan et al., 2018; Oosterhuis et al., 2023; see also the dynamic framework proposed by the STAC-R model Reuter-Lorenz & Park, 2023). Relatedly, in our PLS model (please refer to Figure 3A), the residual variability highlights that age-invariant factors (e.g., lifestyle, leisure activities, socio-demographics) could affect LP performance (see also Gallo et al., 2021 for the association between residual-based measures and cognitive trajectories in late life). Indeed, we observed that the DG component (LC1) was also driven by brain structure changes in white matter networks, showing no significant association with age alone. This further ties in with our hypothesis, stating that DG processes are associated with reserve aging, potentially indexing the age-invariant search and retrieval of less salient semantic knowledge (Hoffman, 2018). Future research should investigate how cognitive reserve factors modulate the reported modified white matter networks, by considering universal and idiosyncratic mechanisms (Baciu et al., 2021).

Similarly, while we controlled for gender differences, our findings would certainly benefit from directly investigating the effect of this variable, especially given that middle-aged women tend to preserve semantic access longer than men as age increases (i.e., higher sentence comprehension performances in late middle age, see Section 2.1.1). In line with this, supplementary analysis suggests that men’s ability to exploit stored semantic resources (LS mechanism) is delayed compared to women. Accordingly, the neurocognitive transition signaling the onset of LP difficulties (i.e., DG-LS interplay) is likely to occur earlier (age 52 vs. 54) (please see Figure B.6, Appendix B for details). This could indicate that all other things being equal, women may more readily manipulate and generalize stored semantic knowledge, contributing to further delay of the onset of word-finding difficulties. Another question remains to understand how this compensatory dynamic pattern translates to pathological aging. For example, one could hypothesize that individuals with neurocognitive disorders would show noticeably limited increases in semantic representations with age or that the ability to access stored semantic knowledge flexibly is compromised early on, resulting in heightened LP deficits. In sum, this perspective offers a promising avenue for establishing whether and how these midlife compensatory mechanisms could capture early signs/trends of neurodegenerative pathology (i.e., neurocognitive biomarkers), paving the way for targeted and anticipated cognitive rehabilitation.

As methodological limitations of our study, we mention that relaxing the orthogonality constraint when performing NMF (Non-Negative Matrix Factorization; see Section 2.3) would be interesting to identify white matter regions overlapping with more than one WM network (Patel et al., 2022, 2023). This is particularly relevant at crossing fiber locations, considering that fiber from different bundle types may cross the same white matter area with a different orientation, thus not necessarily sharing the same component from the NMF solution. Additionally, we acknowledge that the voxel-wise analyses conducted in this study are insufficient for distinguishing fiber differences in white matter alterations in crossing fiber locations. Further analysis showed that cognitive control decline in our voxel-level PLS model was partially associated with a putative integrity increase from the corpus callosum’s splenium extending to the right TPJ (see Figure B.4, Appendix B). Specifically, the bundles converging to the right TPJ (see Figure B.7, Appendix B) have been reported to form a bottleneck region of crossing fibers along an anterior-posterior axis (Schilling et al., 2022). This suggests that this localized increase may be deceitful and instead reflect fiber-specific degeneration (Han et al., 2023). Considering that age-related differences have also been reported in crossing fibers in addition to individual fiber segments (Kelley et al., 2019), fixel-based analysis (FBA) would have been more accurate (Dhollander et al., 2021). However, we preferred not to perform FBA in this study, considering that the b-values (1000 and 2000 s/mm²) in this study are insufficient to attenuate the extra-axonal water signal, meaning that signals from the extra-cellular space outside the axons would contribute to a biased apparent fiber density, rendering “biological interpretation challenging and fundamentally limited” (p.9, Dhollander et al., 2021). Additionally, to our knowledge, MRtrix3 (version 3.0.4; Tournier et al., 2019), does not yet offer users a weighted variant of the streamline-to-voxel algorithm. This means that track-weighted-FA images could not be weighted by the length of the streamline traversing the voxel, resulting in a less accurate contrast.

Finally, out-of-sample replication using larger, independent, and longitudinal databases would also enhance the reliability of our results in light of recent reports showing that cross-sectional studies may largely be underpowered for brain-behavior associations (Gratton et al., 2022; Marek et al., 2022). Additional investigations using a multiple DTI metric (e.g., AD, RD) would also give us a better outlook on age-related biological changes such as demyelination, axonal damage, as well as mild or chronic microstructural alterations (Molloy et al., 2021), leveraging sophisticated machine learning techniques (e.g., Mishra & Liland, 2022)

## 5. Conclusion

In this study, we aimed to disentangle the WM changes related to lexical production (LP) difficulties, typically beginning in middle age. Our findings underscore that midlife is a pivotal period characterized by a discontinuity in brain structure within distributed networks mainly comprised of dorsal, ventral, and anterior cortico-subcortical pathways. Importantly, this discontinuity signals a neurocognitive transition around age 53-54, marking the onset of LP decline. While middle-aged adults may initially adopt a “semantic strategy” to compensate for the initial LP challenges, this strategy may be compromised as late middle-aged adults (age 55-60) lose the ability to exert cognitive control over semantic representations. Taken together, our study (i) emphasizes the importance of considering the interplay between domain-general and language-specific processes when studying the cerebral substrates of lexical production and (ii) reaffirms that sophisticated statistical analysis techniques studies applied to the middle-aged population is a promising avenue for identifying predictive biomarkers of neurodegenerative pathologies.

## Data and code availability statement

Code is available at https://github.com/LPNC-LANG/MiddleAge_LARA2024. Supplementary files for MRI data processing and visualization can be found at https://zenodo.org/doi/10.5281/zenodo.10423906.

## Acknowledgments

This work has been supported by the ANR project ANR-15-IDEX-02. This project has received financial support from the CNRS through the MITI interdisciplinary programs. The Cambridge Centre for Ageing and Neuroscience (Cam-CAN) research was supported by the Biotechnology and Biological Sciences Research Council (grant number BB/H008217/1).

## Conflict of interest statement

Authors declare no conflict of interest.

## Supplementary materials

**Appendix A.** Supplementary material and methods.

**Appendix B.** Supplementary results.

## Notes

https://zenodo.org/doi/10.5281/zenodo.10423907

### Competing Interest Statement

The authors have declared no competing interest.

