## Appendix A for "Dynamics of White Matter Architecture in Lexical Production among Middle-Aged Adults"

**Appendix A: Supplementary material and method**

**Description of the 8 neuropsychological tasks (section 2.1.2)**

**Cattell:** Cattell Culture Fair Test: nonverbal puzzles involving series completion, classification, matrices, and conditions(Cattell & Cattell, 1960).

**Hotel Task:** Simulated tasks of a hotel manager: write customer bills, sort money, proofread adverts, sort playing cards, alphabetize a list of names. Total time must be allocated equally between tasks; there is not enough time to complete any task (Shallice & Burgess, 1991). In this study, we subtracted the log from 1 to stay consistent with a decrease as age increases (i.e., 1-log(x)).

**Naming**: Name the pictured object presented alone (baseline), then when preceded by a prime object that is phonologically related (one, two initial phonemes), semantically related (low, high relatedness), or unrelated (Clarke et al., 2013).

**Proverb:** Read and interpret three English proverbs (Huppert et al., 1994).

**Sentence Comprehension:** Judge grammatical acceptability of partial auditory sentences that begin with an ambiguous sentence stem (e.g., “Tom noticed that landing planes…”) followed by a disambiguating continuation word (e.g., “are”) in a different voice. Ambiguity is either semantic or syntactic, with empirically determined dominant and subordinate interpretations (Rodd et al., 2010).

**Story Recall:** Listen to a short story, recall freely immediately after, then again after a delay, and finally answer recognition memory questions (Tulsky et al., 2003). Delayed recall measure used here.

**Tip-of-the-Tongue:** Participants are asked to name famous faces and indicate if they know/don’t know/or have a ToT (Brown & McNeill, 1966). In this study, we subtracted the score from 1 to stay consistent with a decrease as age increases (i.e., 1-x).

**Verbal Fluency:** Mean of letter (phonemic) fluency and animal (semantic) fluency task. For the phonemic fluency task, participants have 1 minute to generate as many words as possible beginning with the letter ‘p’. For the semantic fluency task, participants have 1 minute to generate as many words as possible in the category “animals” (Lezak et al., 2012).

| Task | Descriptive Statistics  (N=155; Mean, standard deviation) |
| --- | --- |
| **Cattell** | M = 33.1, SD = 5.4 |
| **Hotel Task** (1-log(x)) | M = -4.46, SD = 0.62 |
| **Naming** | M = 0.81, SD = 0.07 |
| **Proverb** | M = 4.87, SD = 1.43 |
| **Sentence Comprehension** | M = 0.90, SD = 0.06 |
| **Story Recall** | M = 13.43, SD = 3.67 |
| **Tip-of-the-Tongue** (1-x) | M = 0.59, SD = 0.02 |
| **Verbal Fluency** | M = 22.01, SD = 5.06 |

**Table A.1. Descriptive statistics of the eight neuropsychological tasks**

**Effect of Age and Gender**

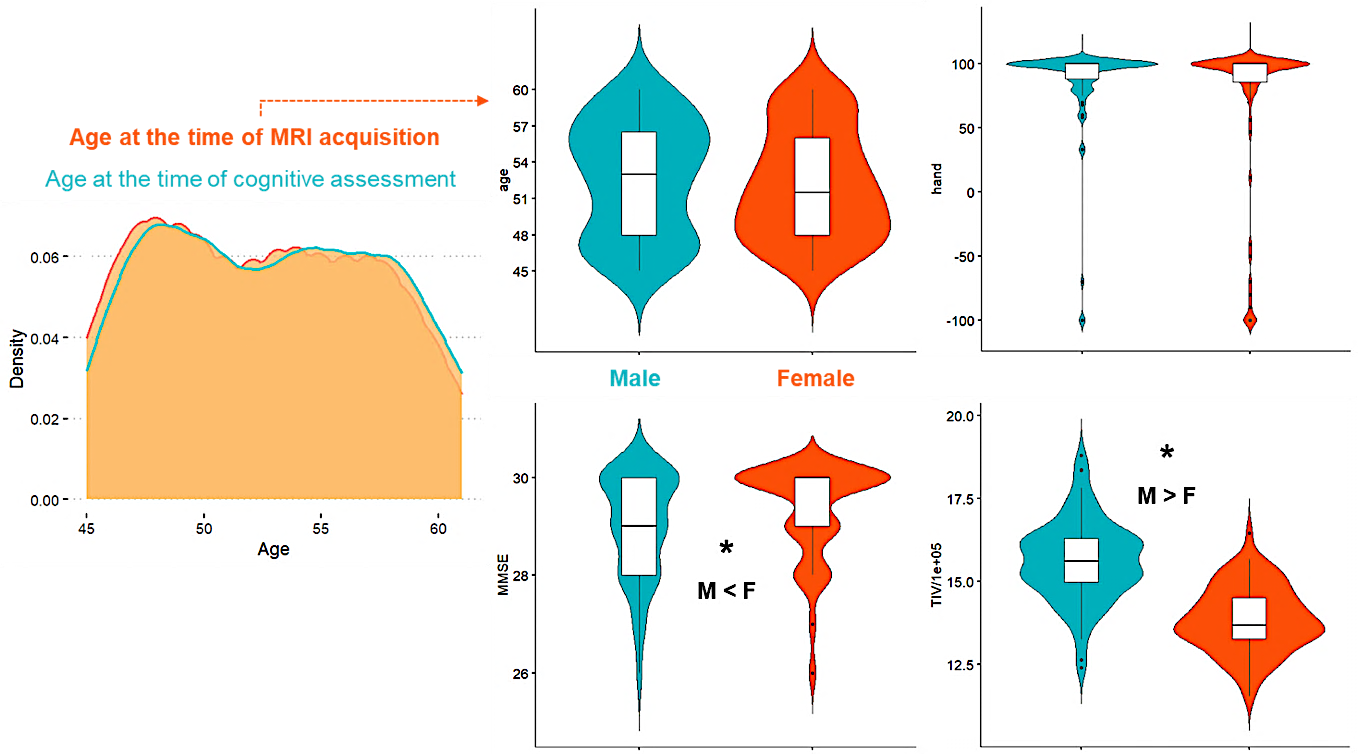
**Figure A.1. Age density plot of the 155-middle-aged-adult sample and gender differences.** When not the response variable, handedness, MMSE score, and TIV were added as covariates along with age at the time of acquisition. Importantly, the distribution of genders across ages was examined without such covariates. Asterisks indicate a significant difference after FDR correction.

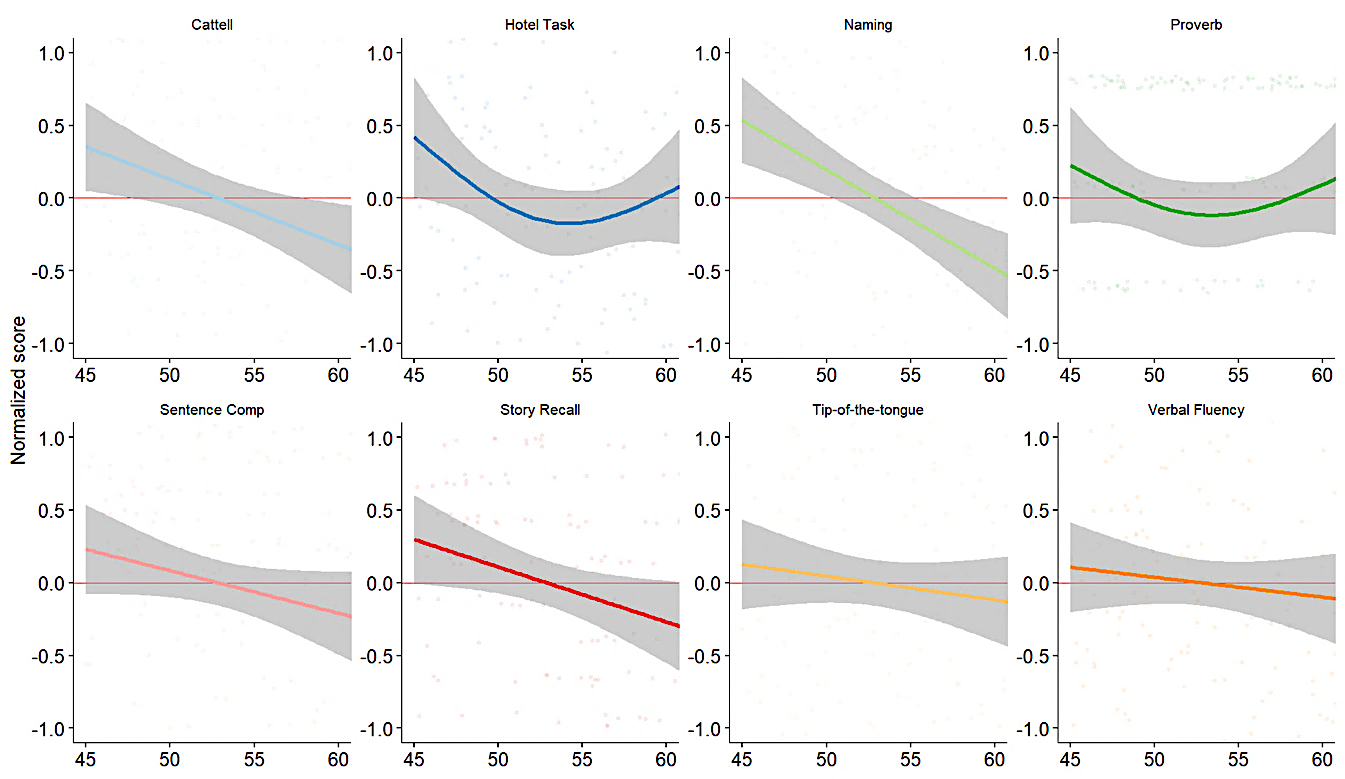

**Figure A.2. Generalized additive models assessing the effect of age across the eight neuropsychological tests during middle age.** We used the age at the time of cognitive assessment for all models. All variables were quantile normalized with the R package *preprocessCore* (v1.60.1; <https://github.com/bmbolstad/preprocessCore>) and scaled for visualization purposes. We used a 3-knot spline to avoid overfitting.

| Neuropsychological tests | Estimate | Std. Error | T value | P-value | FDR-corrected | $\eta_{p}^{2}$ |
| --- | --- | --- | --- | --- | --- | --- |
| Cattell | 0.15 | 1.1 | 0.14 | .89 | .89 | / |
| Proverb | 0.08 | 0.3 | 0.25 | .8 | .89 | / |
| **Naming** | **0.04** | **0.01** | **2.78** | **.006**** | **.04*** | **.04** |
| Tip-of-the-tongue | 0.05 | 0.04 | 1.13 | .26 | .42 | / |
| Hotel Task | 0.1 | 0.13 | 0.78 | .44 | .59 | / |
| **Sentence Comprehension** | **0.03** | **0.01** | **2.61** | **.01*** | **.04*** | **.04** |
| **Story Recall** | **1.6** | **0.7** | **2.34** | **.02*** | **.05~** | **.03** |
| Verbal Fluency | 1.5 | 1.1 | 1.4 | .15 | .32 | / |

**Table A.2. Results of linear models assessing gender differences across neuropsychological tests.** Models are open-sourced and available at <https://github.com/LPNC-LANG/MiddleAge_LARA2024>.

**Preliminary analysis (section 2.3.1)**

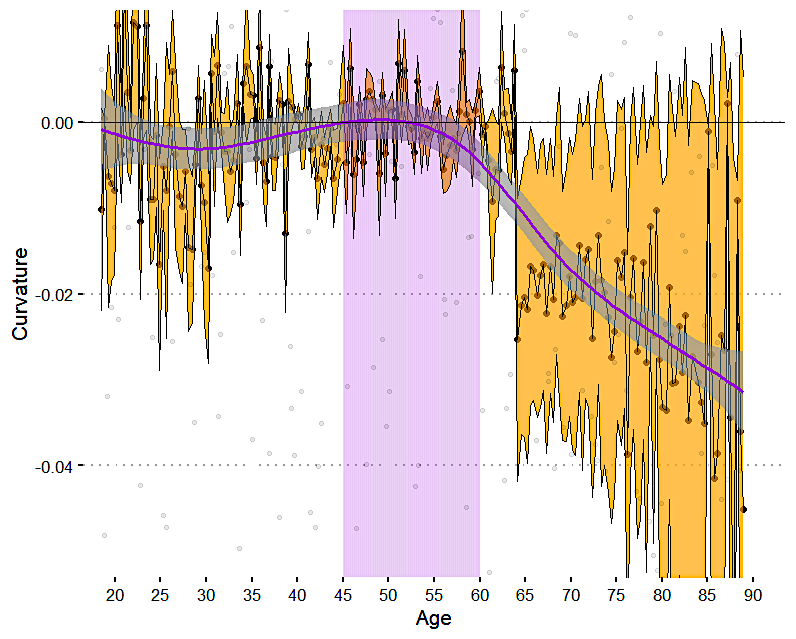

**Figure A.3. Averaged curvature (2^nd^ order derivative) across cognitive variables as age increases.** We used the age at the time of cognitive assessment for all models. To compute curvature, we used the R package *gratia* (v0.8.1; <https://gavinsimpson.github.io/gratia/>; function derivatives with arguments: order = 2, interval = “confidence”, unconditional = “FALSE”). We then selected the significant values for each variable at each time point (i.e., confidence intervals do not contain 0). The inflection point of cognitive aging corresponds to the point at which the curvature crosses zero (i.e., changes its sign). The trajectory was generated for visualization purposes using penalized splines. The ribbon spans the significant confidence intervals' mean lower and upper boundaries.

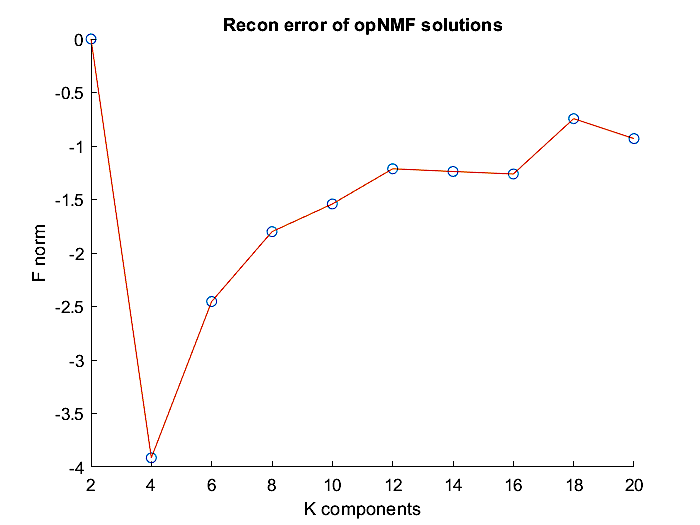
**Neuroanatomical level: Dimensionality reduction (section 2.3.2)***.*

**Figure A.4. Diagnostic plot for finding the optimal number of NMF networks.** Gradient error reconstruction from k=2 to 20 in steps of 2. Please refer to NNMF.m to see the corresponding code used to generate this figure. From k = 16 to 18, a diminishing return on model fit exists. That is, reconstruction error is not minimized as much as before. Following this observation, we choose the 16-network NMF solution.

**Network construction**

From the NMF solution, we build each network using a winner-takes-all approach (cf. NMF_postproc.m at <https://github.com/LPNC-LANG/MiddleAge_LARA2024>). First, we converted the probabilities encoded in the W matrix (*voxels x networks*) into discrete networks by making each voxel take on the column index associated with the highest probability. We then projected these networks onto a nifti volume within the boundaries imposed by the group mask and further filtered this volume to retain clusters of at least 25 connected voxels. The corresponding W and H matrices obtained from the NMF solution can be retrieved at <https://10.5281/zenodo.10423907>.

**Network composition**

To determine the composition of each network, we performed an automatic segmentation of 72 major bundles from the study template SH peaks using TractSeg (Wasserthal et al., 2018) and derived two metrics based on **volumetric** (i.e., spatial overlap at the voxel level) and **connectivity-based information** (at the fiber level, Bullock et al., 2022).

1. Spatial overlap was calculated with the volumetric bundle masks and provided a voxel-wise assessment of the overlap between networks and bundles. Accordingly, we derived a bundle-focused metric (i.e., how much of a given bundle, percentage-wise, overlaps with a given network) and a network-focused metric (i.e., how much of a given network, percentage-wise, overlaps with a given bundle).

2. Connectivity overlap was calculated with the track orientation maps (TOM) and provided a fiber-wise assessment of the overlap between networks and bundles i.e., how many fibers of a given bundle, percentage-wise, transits via a given network

Based on spatial and connectivity information, we created a composite z-scored metric by scaling the 3 metrics before averaging them. Bundles with a composite score above 1 were considered to show a significant contribution to the network. Using this approach, a bundle that would be disregarded based on a limited spatial overlap could be considered as making a noticeable contribution if a high number of its bundles terminated within the limited overlapping region within the network. Conversely, a bundle with high spatial overlap may not necessarily have a high connectivity profile if the overlapping region is not densely populated with the corresponding bundle fibers.
