## Appendix B for "Dynamics of White Matter Architecture in Lexical Production among Middle-Aged Adults"

**Appendix B: Supplementary results**

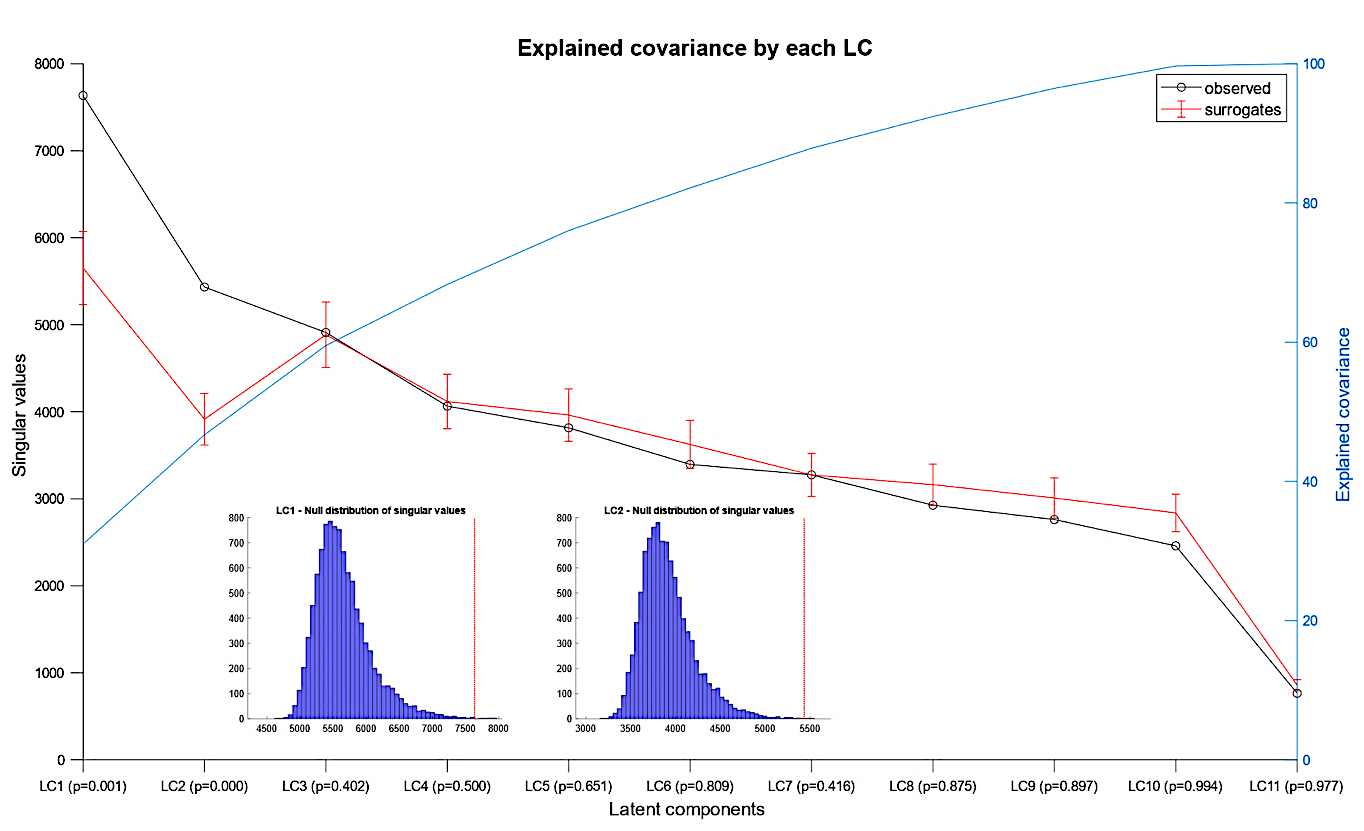
**Figure B.1. Voxel-based PLS diagnostic plots.** Null distribution are based on 10,000 permutations.

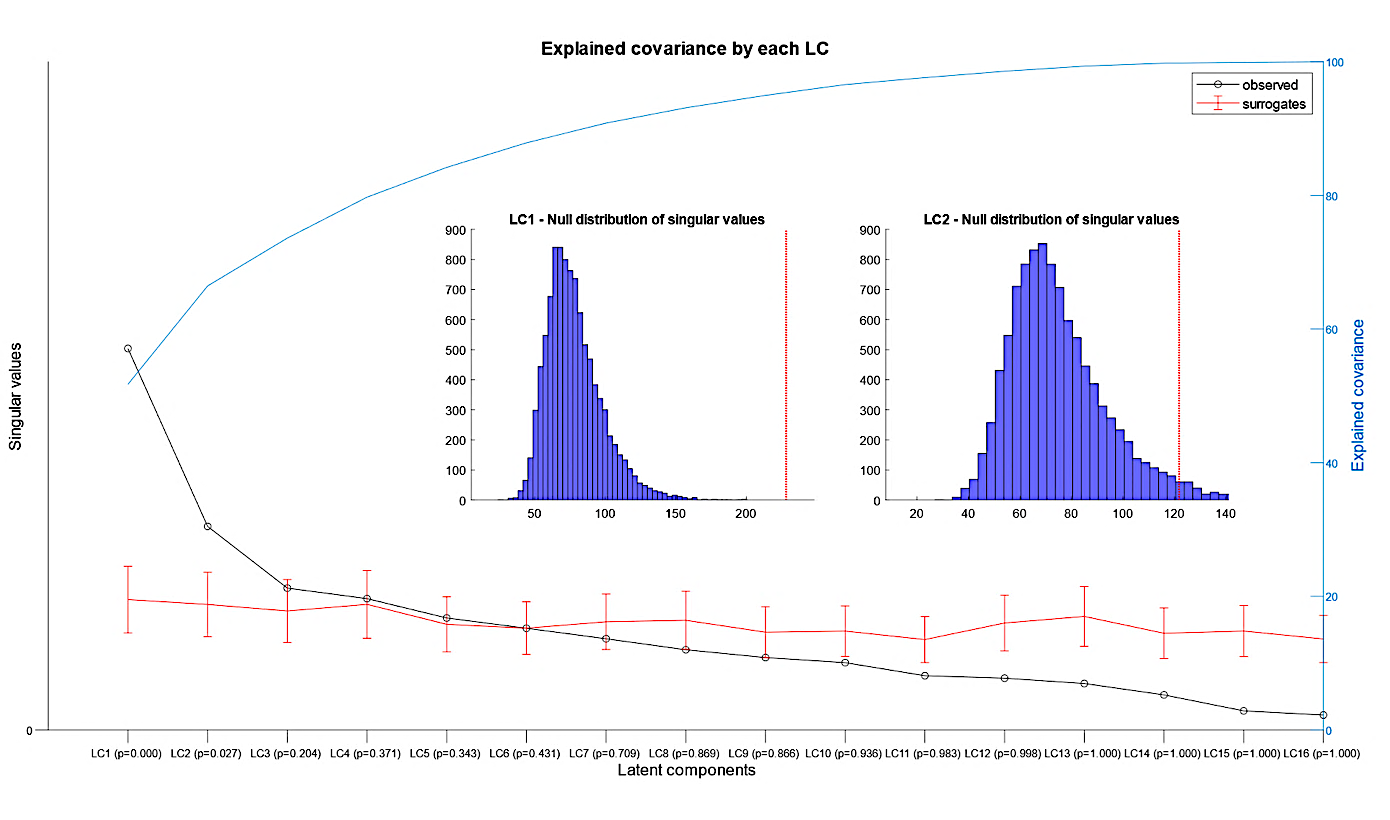
**Figure B.2. Network-based PLS diagnostic plots.** Null distribution are based on 10,000 permutations. LC2 is not significant.

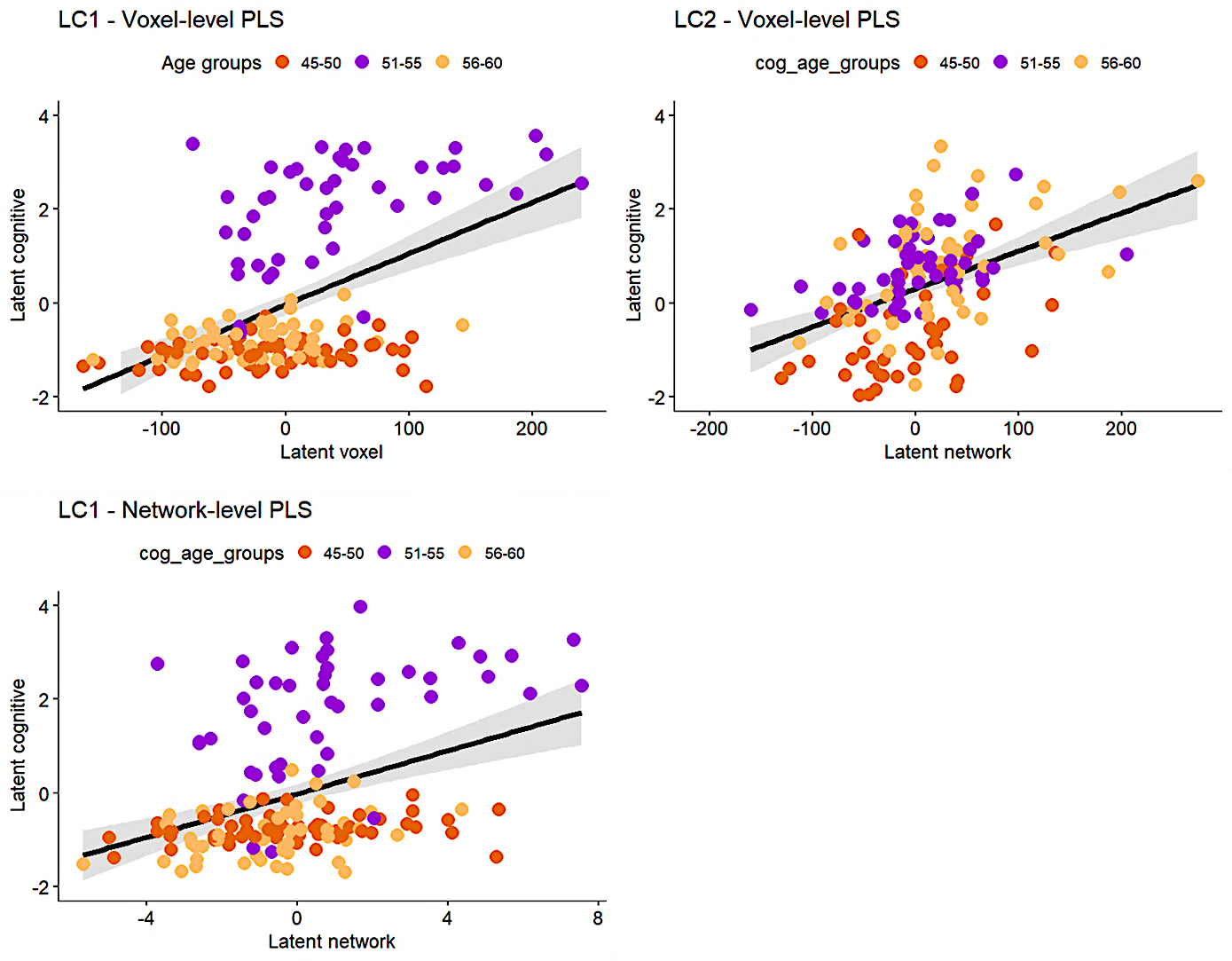
**Figure B.3. Latent scores across age groups.** Figures on the left panel indicates that there is large inter-individual variability in late middle-aged adults in LC1 (domain-general), especially regarding white matter integrity change.

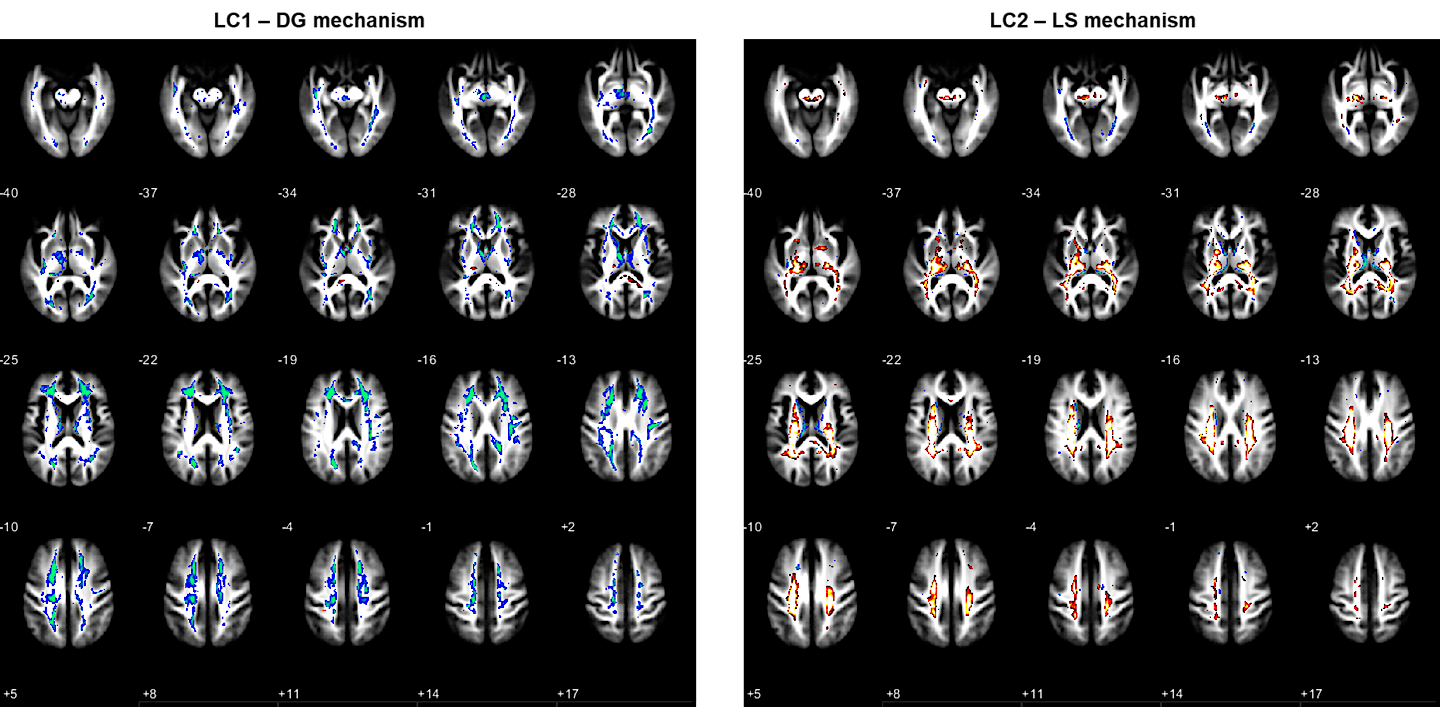

**Figure B.4. Raw brain saliences for LC1 and LC2 in voxel-level PLS**

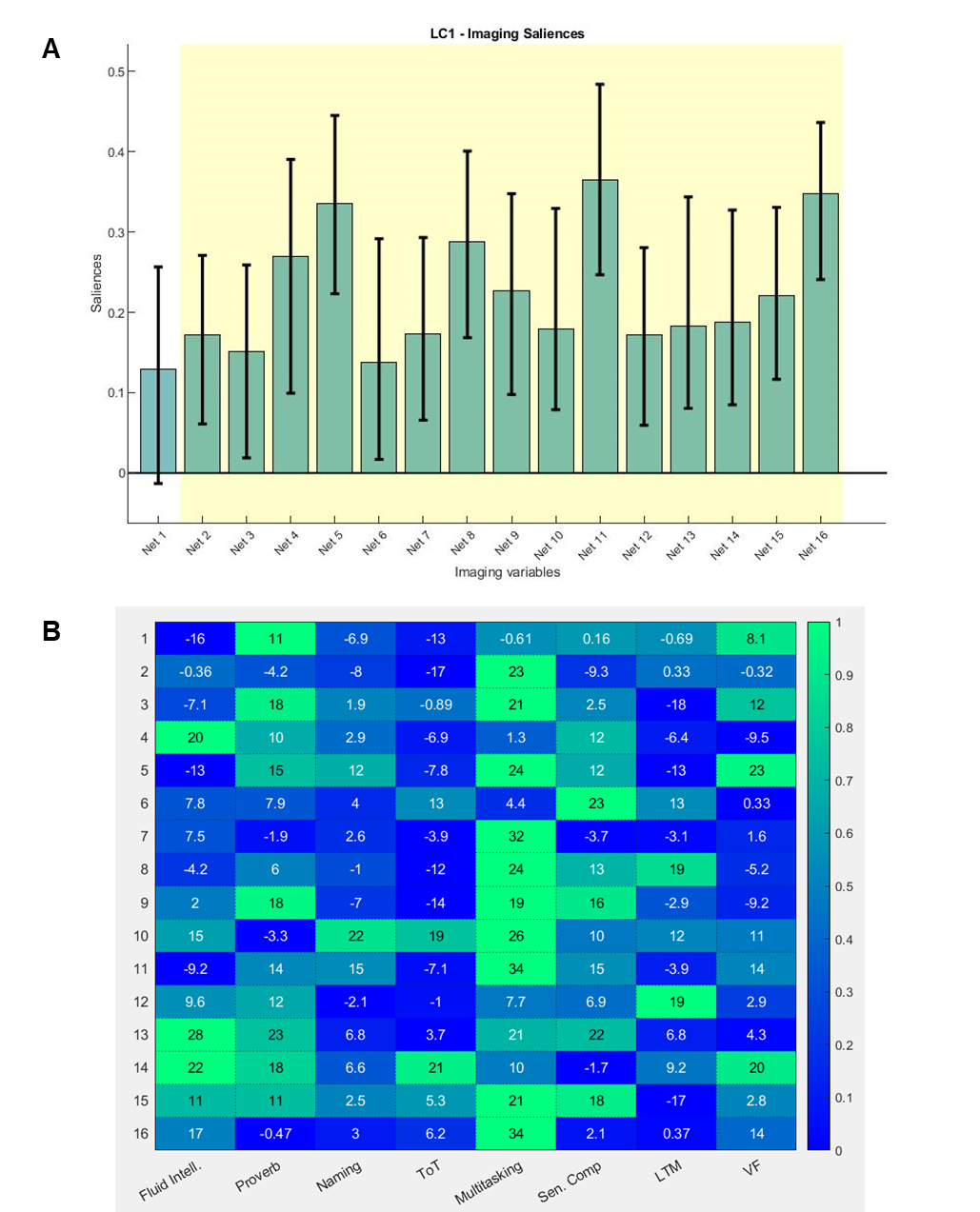

**Figure B.5. Network-level PLS. (A) LC1 Saliences and bootstrapped standard deviations.** Networks highlighted in yellow show robust contribution, suggesting that age-invariant networks also contribute to LP performances. The nifty files associated with each volume can be accessed at <https://10.5281/zenodo.10423907>. **(B) Cross-covariance matrix** between all networks and all cognitive variables.

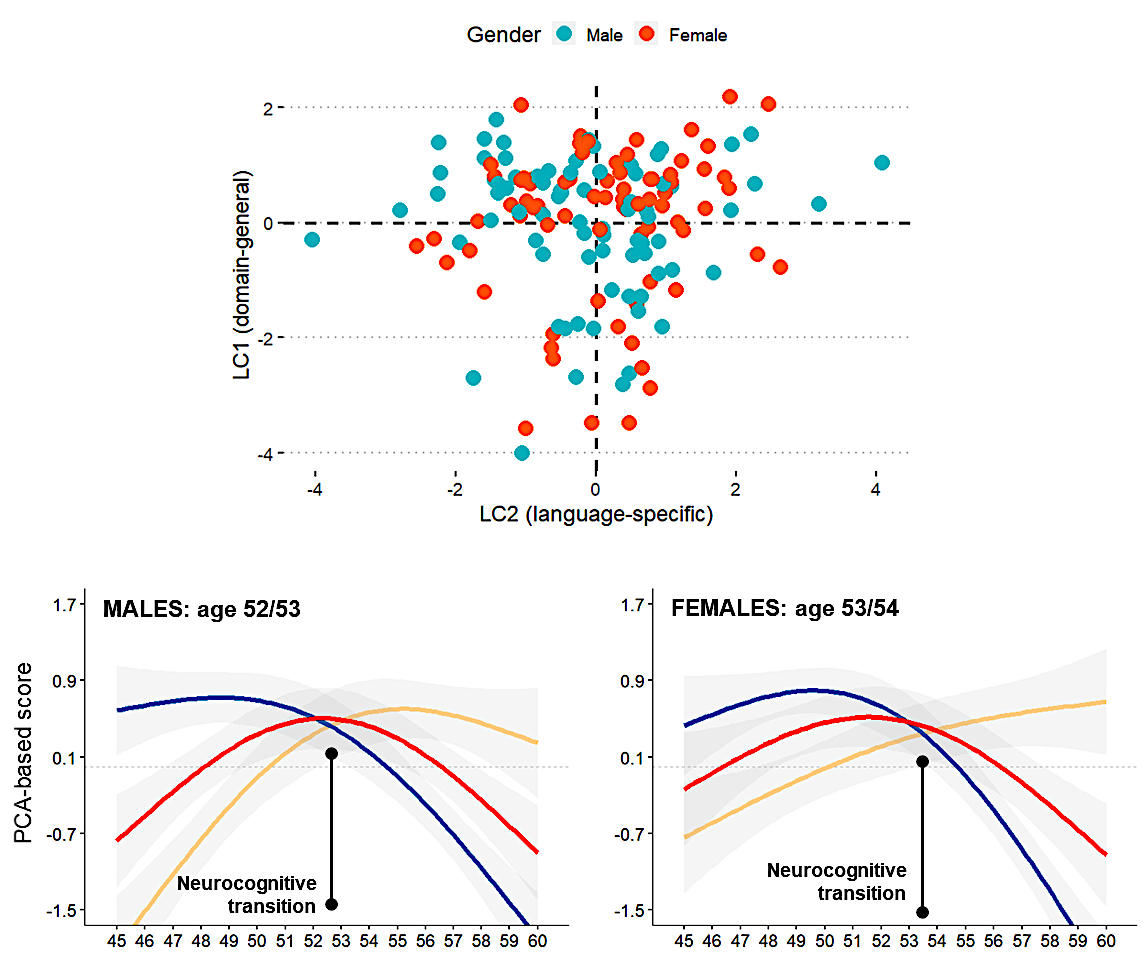

**Figure B.6. Gender effect of the compensatory dynamic of lexical production (LP).** In the bottom panel, blue represents the DG/cognitive control mechanism, yellow represents the LS/semantic mechanism, and red represents their interplay or the control over semantic representations required to maintain LP performance.

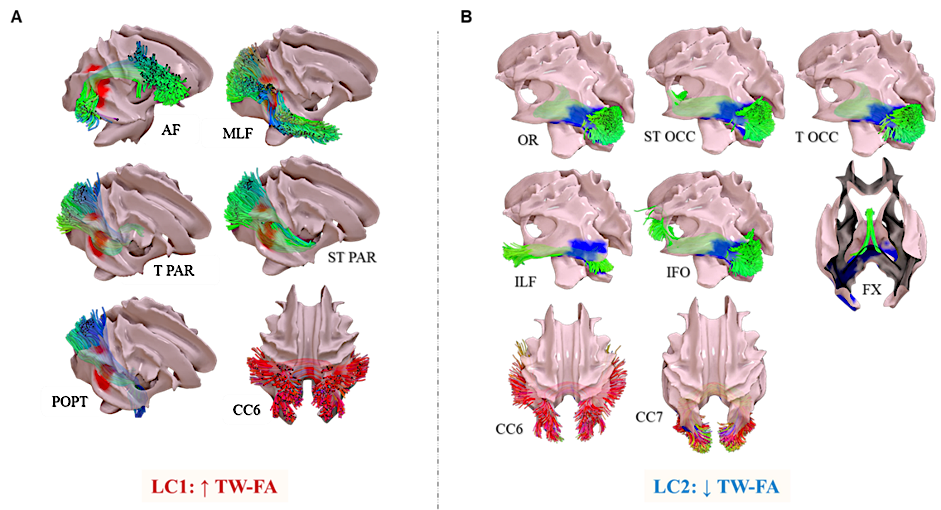

**Figure B.7. Crossing fiber locations.** (A) Illustrates the bundles crossing a region (in red in Figure S4; left panel z = -28) whose increase in TW-FA is paradoxically associated with LP decline (LC1). Tractograms are filtered to retain only the fibers traversing the cluster with a minimum length of 25 mm. Displayed on study template mesh of an averaged TW-FA map. Bundle names correspond to TractSeg’s nomenclature (<https://github.com/MIC-DKFZ/TractSeg>).
